# Dual Ethanolamine Head Groups in Ionizable Lipids Facilitate Phospholipid-free Stable Nanoparticle Formulation for Augmented and Safer mRNA Delivery

**DOI:** 10.1101/2023.10.13.562139

**Authors:** Jing Lu, Ting Jiang, Xuechun Li, Shudan Tan, Yunsen Zhang, Hao Gu, Lin Chen, Junxiang Guo, Rui Yu, Jieying Zang, Defang Ouyang, Hang Yu, Hongwei Yao, Min Qiu, Jinzhong Lin

**Author notes:** These authors contribute equally (Jing Lu, Ting Jiang). Corresponding Authors: (J. Lin); (J. Lu).

## Abstract

The accelerated advent of mRNA-based therapeutics and vaccines, highlighted by the battle against SARS-CoV-2, underscores the urgency to refine lipid nanoparticles (LNPs) for efficient mRNA delivery. In this work, we introduce a novel series of ionizable lipids characterized by double ethanolamine head groups, significantly amplifying mRNA binding affinity. A succinct three-component formulation is subsequently delineated, obviating the conventional dependency on phospholipids inherent in traditional four-component LNPs. Intriguingly, this formulation enables particle formation under neutral pH conditions, a notable departure from the acidic milieu traditionally required, attributable to the enhanced nonionic interactions predominating in mRNA encapsulation. The resultant particles exhibit exceptional stability, superior mRNA encapsulation efficiency, and maintain robust delivery efficacy. When deployed as a vaccine platform, the formulation elicited pronounced humoral and T-cell immune responses, concurrently exhibiting a favorable toxicity profile with a reduced induction of pro-inflammatory cytokines such as IL-6. Our exploration suggests that by fine-tuning the non-electrostatic interactions between the ionizable lipid and mRNA, the dynamics of particle formation can be considerably divergent from the prevailing paradigms of mRNA-LNP formation, hinting at a broader horizon for lipid optimization within the realm of mRNA delivery systems.

## 1. Introduction

The remarkable success of messenger RNA (mRNA) vaccines in combating the threat of severe acute respiratory syndrome coronavirus 2 (SARS-CoV-2) has significantly invigorated the scientific community^1,2^. These mRNA vaccines, known for their robust immune responses, excellent safety profiles, and rapid development pace, have outperformed traditional vaccine approaches^3,4^. However, the success of the mRNA vaccines hinges on the availability of an optimal delivery vehicle considering mRNA’s inherent fragility and poor membrane permeability^5^. Addressing these challenges, lipid nanoparticles (LNPs) have emerged as clinically validated RNA delivery systems, exemplified by the FDA-approved siRNA-LNP drug and two SARS-CoV-2 mRNA vaccines, alongside numerous ongoing clinical trials for LNP-based mRNA therapies^3,6,7^.

Conventional LNP formulations typically comprise four essential components: an ionizable lipid, cholesterol, a “helper” phospholipid, and a PEGylated lipid (PEG-lipid)^8^. Each of these components in LNPs has a crucial role either in mRNA encapsulation or delivery. Among them, the ionizable lipid takes center stage with dual functions. It plays a vital role in determining the encapsulation efficiency of mRNA by electrostatic interactions between the lipid and mRNA. Moreover, this lipid aids in facilitating the escape of cargos from endosome through pH-triggered protonation within these compartments ^9–11^. Extensive efforts have been invested in designing and optimizing the structure of ionizable lipids through traditional combinatorial medicinal chemistry approaches to enable efficient mRNA delivery^12–18^. Notably, recent studies have shed light on the significance of nonionic interactions, such as hydrogen bonds in nucleating the RNA-lipid complex core and enhancing mRNA delivery. For instance, lipids like SM-102 and ALC-0315 used in FDA-approved COVID-19 mRNA vaccines, feature a hydroxyl group attached to the amine linkage.

Apart from the ionizable lipid, cholesterol and PEG-lipid have established roles in LNPs. Cholesterol bolsters LNP stability and promotes RNA transfection, while PEG-lipid regulates particle size^19^, dispersion, prolongs circulation, and prevents aggregation^20^. Phospholipids, such as DSPC (1,2-distearoyl-sn-glycero-3-phosphocholine) and DOPE (1,2-dioleoyl-sn-glycero-3-phosphoethanolamine) often overlooked, are a crucial component influencing the delivery of LNP-mRNA among the four essential components. Phospholipids are the predominant constituents of liposomes employed in drug delivery systems^21^, and their significance becomes evident when developing delivery platforms for nucleic acids, spanning from DNA to siRNA and mRNA. Over time, the proportion of phospholipids has progressively decreased, dwindling from approximately 85 mol% in stabilized plasmid-lipid particles (SPLP)^22^ to 25 mol% in stable antisense lipid particles (SALP) formulations, and further to 20% in stable nucleic acid lipid particles (SNALP) siRNA formulations. In current commercial LNPs designed for siRNA^7^ or mRNA delivery^3^, they contain approximately 10% DSPC. Phospholipids are now designated as “helper” lipid in the LNP system due to its role in nucleic acid encapsulation and stable particle formation, influencing in vivo delivery efficacy. Remarkably, recent research has unveiled that an increased proportion of phospholipids can enhance mRNA delivery efficacy both *in vitro* and *in vivo*, particularly when eff sphingomyelin (ESM) substitutes DSPC^23^. Nonetheless, DSPC tends to favor bilayer structures, resulting in LNP formation with bilayer bleb-like structures under stressful conditions^24^ or when elevated DSPC quantities are employed^25^. Replacing DSPC with DOPE circumvents bleb formation, as DOPE prefers a hexagonal H phase, which is incompatible with bilayer structures^26,27^. In essence, phospholipids are deemed indispensable in LNP formation, with their role extending to encapsulation, particle stability, and overall efficacy in nucleic acid delivery systems.

In this study, our primary objective was to design a novel series of ionizable lipids, with a particular emphasis on the composition of the head group. Through our research efforts, we successfully identified a distinct class of ionizable lipids featuring double ethanolamine head groups. As we delved deeper into the optimization of our formulations, an intriguing revelation emerged: the commonly used “helper” lipid DSPC had a detrimental effect on both mRNA delivery and particle stability. Mechanistic investigations unveiled that these new lipid properties stemmed from the exceptional binding affinity of the ionizable lipid, largely attributed to its unique head group configuration. What was particularly interesting was that the hydrogen-bond interactions overshadowed the contribution of ionic interactions. This unique ionizable lipid could efficiently encapsulate mRNA without the necessitating use of an acidic buffer. Moreover, it rendered phospholipids unnecessary for achieving stable LNP formation and long-term storage, all while maintaining high mRNA encapsulation efficiency.

Our research has culminated in the formulation of an ultra-stable, phospholipid-free, three-component nanoparticle architecture for mRNA encapsulation, designated as the iPLX formulation to distinguish it from the conventional four-component LNP formulation. This novel formulation, particularly with the inclusion of the leading compound LQ-3, exhibited substantial potential. When deployed as a vaccine platform, LQ-3 iPLX elicited robust humoral and T-cell immune responses. Furthermore, LQ-3 iPLX manifested promising toxicity profiles, notably inducing a reduced quantity of pro-inflammatory cytokines such as IL-6 and TNF-α.

Our study not only points to a new direction for optimizing ionizable lipids but also underscores the pivotal role of tuning the interaction between the head group and nucleic acid. This allows for the creation of stable particles using ionizable lipids, cholesterol, and PEG-lipids, with remarkable mRNA encapsulation efficiency while maintaining superior delivery efficacy.

## 2. Results

### 2.1. Synthesis, Preparation, and *In Vivo* Screening of Ionizable Lipids with Double Ethanolamine Head Groups

At the heart of effective LNP-mediated mRNA delivery lies the role of ionizable lipids, orchestrating mRNA binding during nanoparticle assembly and facilitating escape from endosomes ^28^. With a strategic focus on enhancing LNP stability, mRNA delivery efficiency, and safety, we embarked on the design of a novel library of chemically distinct ionizable lipids. The significance of tertiary amines that acquire a positive charge upon protonation in a low pH environment has been extensively explored, facilitating their interaction with the negatively-charged RNA molecules^29^. Recent advancements have brought attention to the potential benefits of incorporating a hydroxyl group onto a tertiary amine to form the lipid headgroups. This augmentation further enhances *in vivo* mRNA delivery of formulated LNP. This advantageous effect has been attributed to the formation of hydrogen bonds between the introduced hydroxyl group and the RNA molecules^14,30^. Drawing inspiration from by these pivotal design cues, we meticulously crafted 14 novel lipids, all featuring dual hydroxyls and tertiary amines within their lipid headgroups. Detailed synthetic routes and structural confirmation by ^1^H NMR were provided in Figures S1-S14 in the Supporting Information. We next prepared LNP-mRNAs by formulating the synthetic ionizable lipid with cholesterol, DSPC, DMG-PEG2k, and firefly luciferase (Fluc) mRNA through a microfluidic system. The lipid composition adhered to a generic mole ratio of 50:10:38.5:1.5 for ionizable lipid, DSPC, cholesterol, and DMG-PEG2k. The structures of the 14 ionizable lipids and the physicochemical attributes of mRNA-encapsulated LNPs (LNPs-mRNA) were presented in Figures 1A and 1B. The resulting mRNA-LNPs showed average particle sizes ranging from of 70 to100 nm, accompanied by low polydispersity indices (PDI) (∼0.1), and mRNA encapsulation efficiency exceeding 96%.

**Figure 1.**
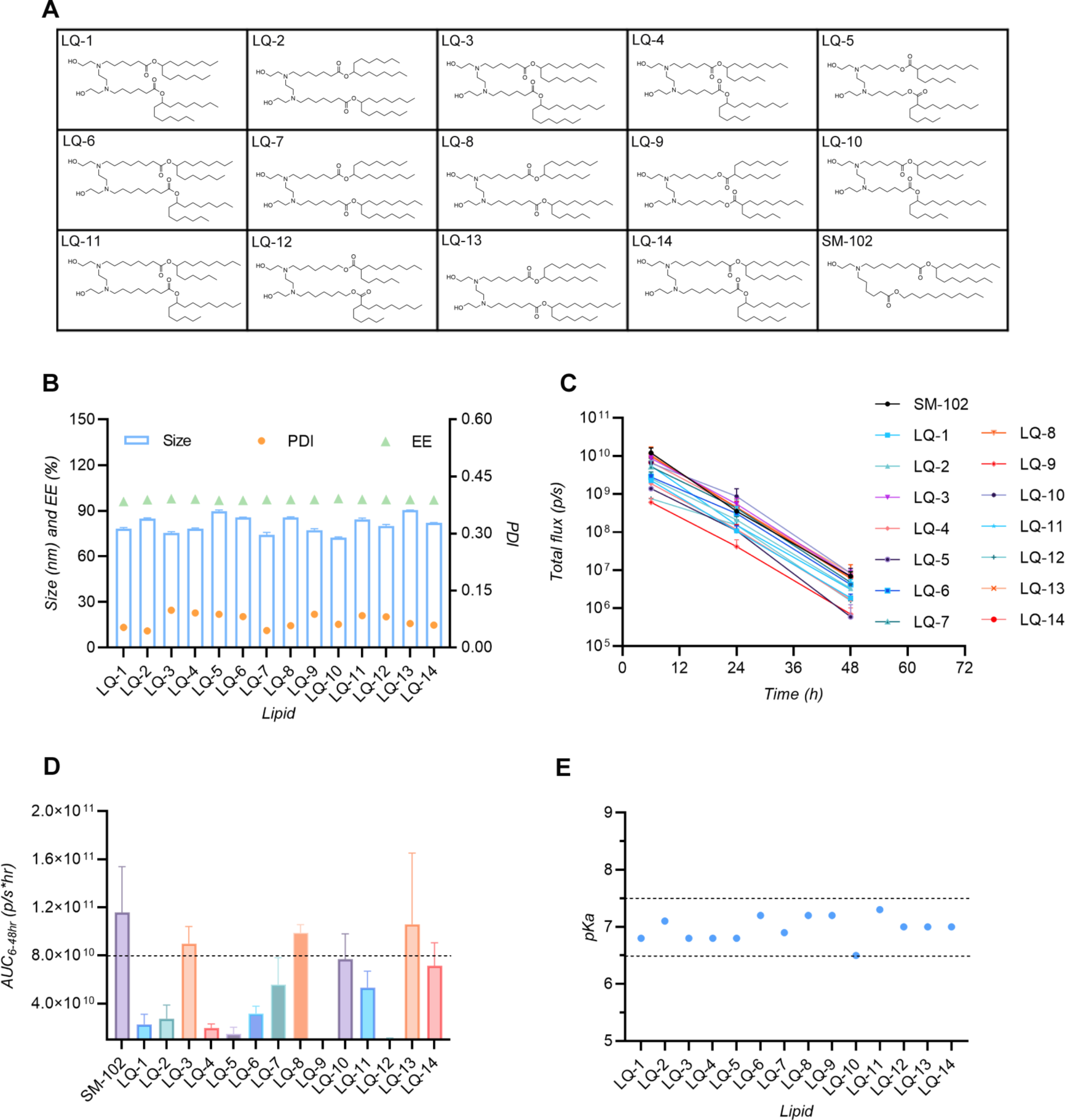
Chemical structures, physicochemical properties, and *in vivo* screening. (A) Chemical structures of synthesized ionizable lipids. (B) Physicochemical properties of LNPs, formulated with a mole% ratio of Ionizable lipid: DSPC: Cholesterol: DMG-PEG2k = 40: 11: 47.4: 1.6, and N/P ratio of 8:1 (n = 3, mean + SD). (C) Luciferase intensities of mRNA-LNPs incorporated with various ionizable lipids in livers at intervals of 6, 24, and 48 hours post i.v. administration at a dose of 0.25 mg/kg mRNA (n = 3, mean + SD). (D) The areas under the curve (AUC) of whole-body luciferase bioluminescent signals for novel LNPs in comparison to SM-102 LNP (n = 3, mean + SD). (E) pKa values of LNPs as measured by TNS.

To explore the impact of these novel ionizable lipids on *in vivo* mRNA delivery efficacy, mRNA-LNPs were intravenously (*i.v.*) administrated into female wild-type Balb/c mice at a dose of 0.25 mg/kg Fluc mRNA, and the structure-activity relationships were explored. Our benchmark for comparison was SM-102, a synthetic lipid utilized in the FDA-approved SARS-CoV-2 mRNA vaccine. Tracking the whole-body bioluminescence intensities of these administered mice at various time points using an *in vivo* imaging system (IVIS) provided insights into the dynamic kinetic profiles of luciferase expression (Figure 1C). Notably, a majority of our 14 LNPs exhibited strikingly similar profiles to that of SM-102 LNP. Interestingly, the expression of mRNA seemed to diminish at a slower rate within the 4 - 24 hour interval for most of these LNPs. As a result, despite SM-102 exhibiting a more pronounced bioluminescence intensity compared to our LNPs at the 4-hour mark post-administration, the cumulative protein expression of our LQ-3, LQ-8, and LQ-13 LNPs closely matched that of SM-102 LNP. This was evident through measurement of the areas under the whole-body luciferase bioluminescent signal curve (AUC) (Figure 1D).

Moreover, it has been well-established that the apparent *p*Ka of LNPs plays an important role in shaping the in *vivo* RNA delivery potency. The optimal *p*Ka range for liver-targeting LNP-mediated RNA delivery resides between 6.0 and 7.0 ^16,31,32^. In line with these established findings, the most potent LNPs, LQ-3 and LQ-13, had a surface *p*Ka of 6.8 and 7.0, respectively (Figure 1E).

### 2.2. Optimizing Nanoparticle Formulation with LQ-3, a Double Ethanolamine-Headed Ionizable Lipid

Following the initial screening phase, LQ-3 emerges as the leading compound, prompting a comprehensive three-round optimization process for its LNP formation. Typically, LNP is a four-component system encompassing an ionizable lipid, phospholipid, cholesterol, and PEG-lipid^8^. Phospholipid, such as DSPC, are known to play a crucial role in encapsulating nucleic acids and providing stability ^27,33^. In our first step, we maintained a molar ratio of ionizable lipid and DMG-PEG2k at 40% and 1.6%, respectively. The proportion of DSPC were adjusted from 0% to 40%, with corresponding variations in cholesterol ratios to maintain a total component ratio of 100%. Luciferase mRNAs were encapsulated at an N:P ratio of 8:1. The designed LNP formulations and the relevant characteristics are presented in Figure 2A and in Table S1 in the Supporting Information. Surprisingly, as DSPC content decreased, the mRNA delivery efficacy of resulting LNPs progressively increased, reaching its peak when zero DSPC was included. Figure 2B illustrates the *in vivo* whole-body bioluminescence intensities 4 hours post-administration of LNPs, while Figure 2C displays the AUC over 4 to 48 hours. When employing DOPE as a substitute for DSPC, a similar trend was observed; the delivery efficiency peaked when the DOPE content was reduced to 2% or entirely omitted (Figure S15).

**Figure 2.**
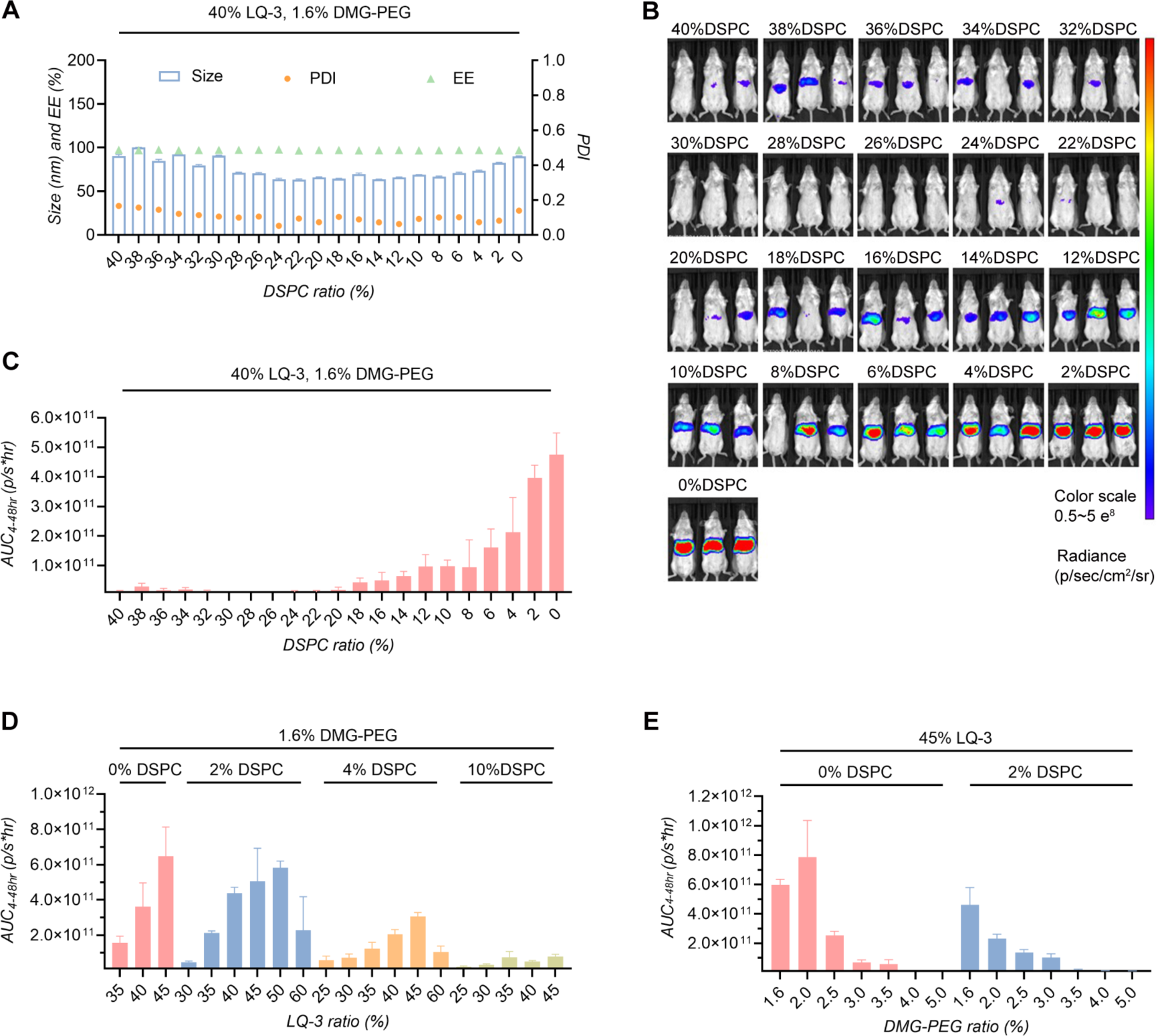
Optimization of LQ-3 nanoparticle formulations for mRNA delivery. (A) Properties of nanoparticles over a wide range of DSPC ratio (n = 3, mean ± SD). (B) Whole-body bioluminescence imaging taken 4 hours after intravenous administration of these nanoparticles at an mRNA dosage of 0.25 mg/kg. (C) Area under the curve (AUC) for liver bioluminescent signals spanning from 4 to 24 hours. (D) Identification of optimal LQ-3 ratios. (E) Determination of the best PEG-lipid ratio.

At the core of LNP dynamics lies the role of the ionizable lipid, influencing both mRNA encapsulation and endosomal escape. In our second optimization, we investigated the effects of ionizable lipid ratio on mRNA delivery. This was achieved through a systematic exploration of 20 formulations with varying LQ-3 content, ranging from 25% to 60%, coupled with 0%, 2%, 4%, and 10% DSPC. Detailed parameters for these LNPs are presented in Table S2 in the Supporting Information. Echoing the outcomes of the first round, LNPs containing 4% and 10% DSPC exhibited notably reduced mRNA delivery efficacy compared to those with 0% and 2 % DSPC. Among the spectrum, LNPs with 40-50% ionizable lipid content demonstrated elevated mRNA delivery efficacy. Particularly noteworthy was the LNP configuration consisting of 45% LQ-3, 0% DSPC, 53.4%, cholesterol, and 1.6% DMG-PEG2k, demonstrating the peak of efficacy (Figure 2D).

Finally, we investigated the impact of PEG-lipid ratio on luciferase expression *in vivo*. Parameters detailing this PEG-lipid-focused screen can be found in Table S3. While maintaining the ionizable lipid/DSPC ratio at 45/0 or 45/2, we examined the expression efficacy of LNPs formulated with different PEG-lipid ratios, ranging from 1.6% to 5%, *in vivo*. The results showed that the formulation with 2% DMG-PEG2k ratio exhibited the highest expression, followed by the 1.6% DMG-PEG2k ratio (Figure 2E). In sum, these results highlight the optimal LQ-3 nanoparticle formulation comprising 45% ionizable lipids, 0% DSPC, 53% cholesterol, and 2% DMG-PEG2k after a three-round optimization process.

### 2.3. Characterization of Phospholipid-Free LQ-3 iPLX Nanoparticles

Our previous investigations into nanoparticle formulations yielded an unexpected discovery: the omission of DSPC, a commonly used “helper” lipid, surprisingly enhanced the efficacy of mRNA delivery, challenging the conventional role of DSPC in LNPs featuring ionizable lipids. While in the realm of lipoplex formulation, where nanoparticles are created through the interaction of cationic lipids and nucleic acids facilitated by cholesterol, the role of DSPC becomes dispensable^34^. In contrast, most ionizable lipid-based formulations have historically relied on DSPC for ensuring stable LNP formation and optimal delivery outcomes.

This intriguing revelation, with our lipid LQ-3 as a prominent example, prompted us to conduct further investigations into this DSPC-free, three-component formulation. To clearly delineate this new approach from conventional formulations, we introduced the term “iPLX” for the three-component variant, drawing parallels with conventional lipoplex formations but integrating an ionizable lipid in place of a cationic lipid, which stands in contrast to the conventional four-component LNP. To ensure robust comparisons and controls, we extended our study by employing SM-102 to create nanoparticles using both iPLX and LNP formulations, providing a comprehensive understanding of our newly identified ionizable lipids within the context of the iPLX formulation.

Both LQ-3 and SM-102 demonstrated the ability to form nanoparticles in the absence of DSPC without a notable decrease in encapsulation efficiency (Figure 3A). Interestingly, transitioning from LNP to iPLX formulation, resulted in a slight size increase for LQ-3 nanoparticles (less than 10 nm), whearas SM-102 iPLX displayed a more significant expansion from 80 nm to 120 nm. Notably, the influence of DSPC on the PDI varied for LQ-3 and SM-102 nanoparticles. For SM-102, the inclusion of DSPC led to a smaller PDI, whereas for LQ-3, the PDI decreased as DSPC was excluded. These findings highlighted the divergent effects of DSPC on LNP formation for LQ-3 and SM-102.

**Figure 3.**
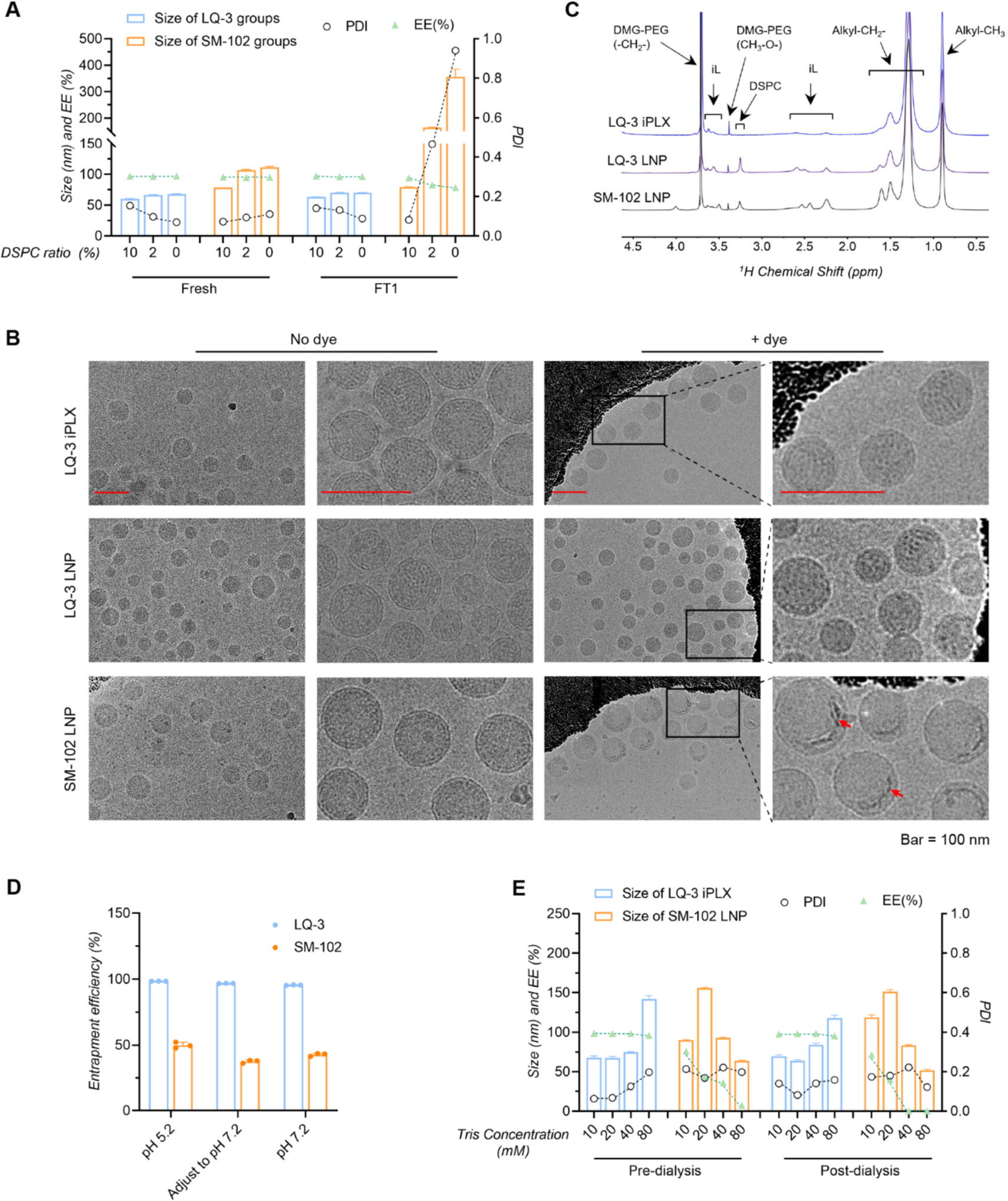
Mechanistic investigation of phospholipid-free LQ-3 iPLX formulation. (A) Evaluation of size, polydispersity index (PDI), and encapsulation efficiency (EE) for LQ-3 and SM-102 nanoparticles across DSPC ratios of 10%, 2%, and 0%. (B) Cryo-EM visualization of LQ-3 iPLX (0% DSPC), LQ-3 LNP (10% DSPC), and SM-102 LNP. Red arrows indicate location of mRNA. Scale bar set at 100 nm. (C) 1H NMR spectra for LQ-3 iPLX, LQ-3 LNP, and SM-102 LNP with “iL” denoting ionizable lipid. (D) Assessment of mRNA binding efficiency for LQ-3 using the RiboGreen assay under acidic and neutral pH conditions, with SM-102 as the reference. (E) Characterization of LQ-3 iPLX and SM-102 LNP formulated with mRNA prepared in Tris-HCl buffer (pH 7.2) at different tris concentrations.

To validate these observations, we subjected these nanoparticles to a freeze-thaw cycle. Intriguingly, SM-102 iPLX nanoparticles underwent collapse, as evidenced by the PDI increasing to nearly 1.0 and the size expanding to 350 nm, accompanied by mRNA release (Figure 3A). Even in the presence of 2% DSPC, SM-102 LNP underwent decomposition, albeit to a lesser extent. Only with 10% DSPC did SM-102 LNP withstand freeze-thaw conditions without substantial changes in particle properties. This underscores the crucial role of DSPC in stabilizing nanoparticles, particularly for ionizable lipid SM-102. This trend likely extends to other well-characterized lipids such as DLIN-MC3 and ALC-0315, both incorporating around 10% DSPC in LNP formulation. In a contrasting scenario, transitioning from LNP to iPLX formulation did not yield noticeable particle alterations for LQ-3 lipid, implying comparable stability between LQ-3 iPLX and LQ-3 LNP. This stability underscores the potential of LQ-3 iPLX as a promising platform for mRNA delivery.

Further exploration through cryoelectron microscopy (Cryo-EM) unveiled the morphological aspects. Both LQ-3 iPLX and LNP showed a spherical structure with a dense core and monolayer shell, akin to SM-102 LNPs (Figure 3B). However, a closer inspection revealed that both LQ-3 iPLX and LNP exhibited a degree of multilamellarity, while SM-102 LNP displayed an amorphous core. To gain further insights, we employed thioine dye staining to visualize the localization of RNA within LNPs using Cryo-EM^35^. In the case of SM-102 LNP, the RNA density was clearly visible on the periphery of the dense core beneath the outer shell, consistent with prior observations, suggesting that ionizable lipids within SM-102 LNP redistributed into the core when equilibrated in a neutral buffer^24^. Interestingly, LQ-3 iPLX and LNP did not exhibit enhanced contrast due to RNA staining, indicating that LQ-3 forms a tighter complex with RNA, impeding the interaction with thionine dye.

Additionally, we harnessed ^1^H NMR spectra to gain further insights into lipid surface distributions within these nanoparticles. Figure 3C reveals distinct hydrogen peaks corresponding to ionizable lipids and PEG-lipids, detected on the surface of LQ-3 iPLX, LQ-3 LNP, and SM-102 LNP. As anticipated, the choline signal from DSPC was absent in the LQ-3 iPLX spectrum, while evident in both the SM-102 and LQ-3 LNP spectra. Consequently, iPLX nanoparticles exhibit a higher content of ionizable lipid on the surface which may contribute to its enhanced mRNA delivery efficacy.

We measured the zeta potential of both LQ-3 iPLX and SM102 LNP in Tris Buffer (pH 7.5). The zeta potential for SM102 LNP is approximately −4 mV, whereas that for LQ-3 iPLX is slightly higher at −2.6 mV (Table S4). This aligns with the observation that the pKa of LQ-3 iPLX is marginally higher than that of SM102 LNP.

### 2.4. Nonionic Interactions as Key Drivers of RNA-Lipid Nucleation in LQ-3 iPLX nanoparticles

The contrasting behaviors exhibited by LQ-3 and SM-102 prompted us to further explore their underlying mechanisms. The role of an ionizable lipid in facilitating the interactions with negatively charged nucleic acids is crucial for their encapsulation and stabilization within lipid-based delivery systems. The primary distinction between LQ-3 and SM-102 lies in the structure LQ-3’s head group, which features double ethanolamine linked by an ethyl bond. Consequently, under acidic conditions, LQ-3’s head group carries two positive charges, enabling ionic interactions with nucleic acids. However, this unique head configuration also amplifies the potential for other electrostatic interactions, such as hydrogen bonding, with nucleic acids. There, it is essential to investegate the role of these interactions in shaping LQ-3’s distinct properties.

We first assessed the capacity of LQ-3 and SM-102 to form complexes with nucleic acids. RNA was dissolved in either an acidic acetate buffer (pH 5.2) or a neutral Tris-HCl buffer (pH 7.2), while LQ-3 and SM-102 were dissolved in ethanol. The lipid solution was then introduced into the mRNA solution under stirring conditions, and we measured encapsulation efficiency using the ribogreen assay. In acidic conditions, LQ-3 exhibited robust mRNA encapsulation, achieving an efficiency of 98% at an N:P ratio of 6:1(Figure 3D). In contrast, only 50% of the mRNA was accessible to ribogreen dye when using SM-102. Remarkably, upon adjusting the pH of the lipid:RNA solution to 7.2, LQ-3 maintained its mRNA complexing ability at an efficiency of 97%, whereas SM-102’s efficiency further decreased to 38%. When the mixing was carried out in a neutral buffer at pH 7.2, LQ-3 effectively encapsulated mRNA through complex formation with 95% protection, while SM-102 only provided 43% mRNA protection.

Building on above findings, we hypothesized that mRNA may not necessarily require an acidic buffer for successful nanoparticle formation. To explore this, we dissolved mRNA in a neutral Tris-HCl buffer (pH 7.2) and rapidly mixed it with lipid mixtures in ethanol using a microfluidic device, following standard procedures. The resulting solution was then dialyzed in the same buffer used for mRNA dissolution to remove ethanol. Considering that ionic interactions can be sensitive to ionic strength, we conducted tests with varying concentrations of Tris-HCl buffer, ranging from 10 mM to 80 mM. Interestingly, our lipid LQ-3, when combined with cholesterol and PEG-lipid, was able to encapsulate mRNA even in a neutral buffer (Figure 3E). Remarkably, even with a higher 80mM Tris concentration, although the particle size expanded to 140 nm, a stable nanoparticle assembly with a 96% encapsulation efficiency was still achievable. These findings hint that the primary forces propelling RNA:lipid complex formation remain unaffected by ionic strength, underscoring the crucial role of nonionic interactions between mRNA and ionizable lipids. In contrast, when utilizing the lipid SM-102, stable nanoparticles could not be formed when mRNA was dissolved in neutral buffer, as initially expected. Additionally, the assembly of SM-102-based nanoparticles was significantly affected by ionic strength. With 10 mM Tris, the encapsulation efficiency was approximately 70%, but this efficiency plummeted to zero when Tris concentrations of 40 mM or higher were used (Figure 3E).

### 2.5. Molecular Dynamics Simulation Reveals Dominance of Nonionic Interactions in mRNA Complexation by LQ-3

The previous data compellingly demonstrates that LQ-3 has a superior propensity to form complexes with mRNA compared to SM-102, possibly due to differences in their headgroups. To discern if nonionic interactions play a part in these disparities, we executed Molecular Dynamics (MD) simulations on both lipids, ensuring the amine groups were deprotonated to exclude ionic interactions. Figures 4A and 4B chart the binding trajectory of lipids SM-102 and LQ-3 with mRNA over a span of 100,000 ps at distinct intervals: 0, 4,000, 15,000, and 100,000 ps, illustrating the lipids’ transition from a free to a bound state. In both scenarios, single-stranded mRNA was observed dynamically enveloping a lipid micelle, becoming apparent around 15,000 ps, with the lipid headgroups engaging with the mRNA nucleobases or phosphate backbone.

**Figure 4.**
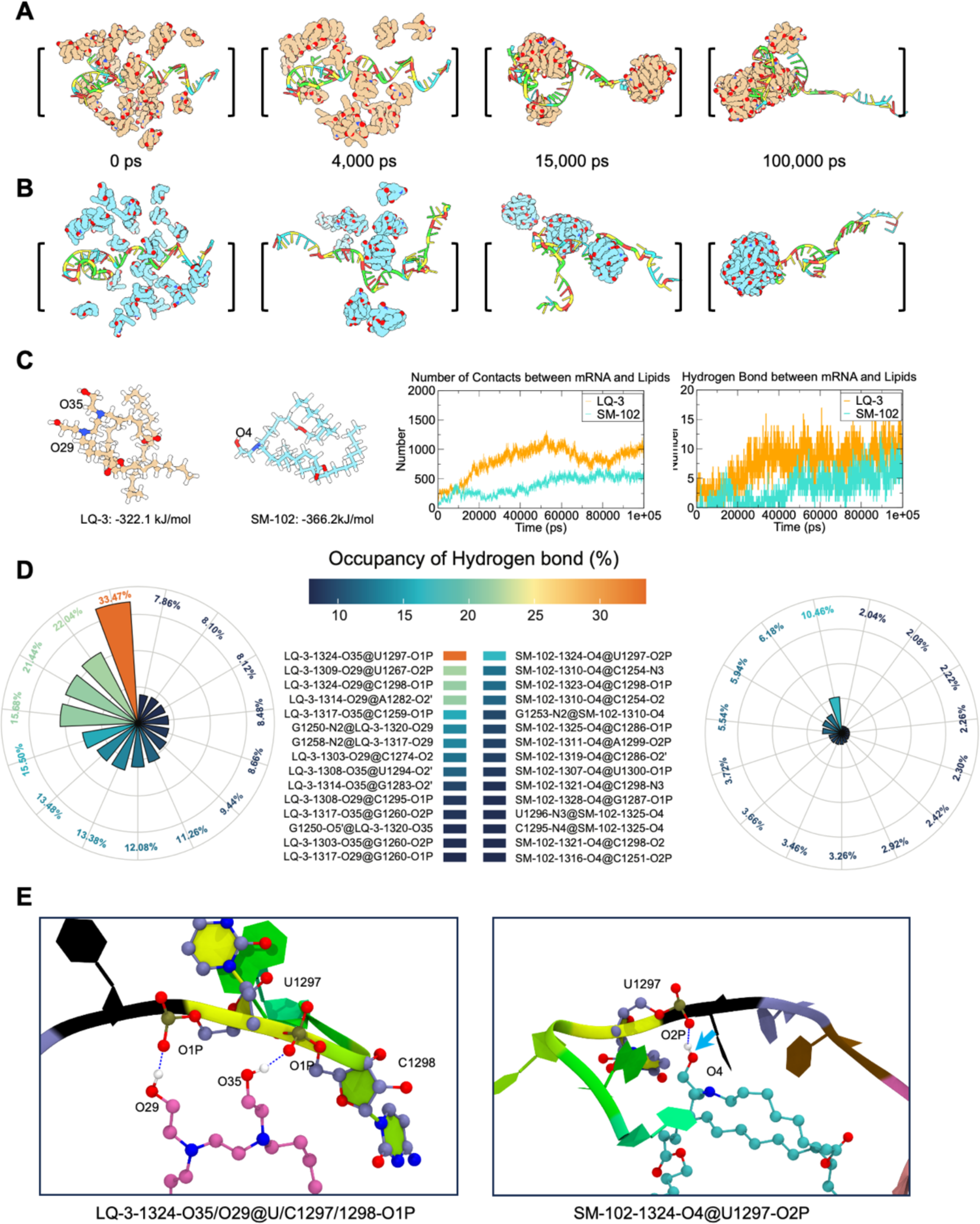
Molecular dynamics simulation. (A-B) Simulation snapshots for LQ-3 (A) and SM-102 (B) captured at 0 ps, 4,000 ps, 15,000 ps, and 100,000 ps. (C) Structures of the two lipids alongside a summary of the total contacts and hydrogen bonds formed between mRNA and the lipids. (D) Nightingale rose charts of the occupancy of the top 15 hydrogen bonds between mRNA and LQ-3 (left) or SM-102 (right). (E) Representative interactions established between mRNA and lipids.

The interaction statistics, including contacts and hydrogen bonds between mRNA and both ionizable lipids, are depicted in Figure 4C. Here, LQ-3 establishes more contacts and hydrogen bonds with mRNA than SM-102, suggesting a stronger interaction affinity for LQ-3 lipid with mRNA compared to SM-102. Correspondingly, interaction energy assessments, outlined in Table S5, unveil a more robust interaction energy for LQ-3 lipid as opposed to SM-102. Specifically, LQ-3 displayed stronger interaction energy (−1236.9 kJ/mol) than SM-102 (−593.4 kJ/mol), with a delta E of - 643.5 kJ/mol. Furthermore, the van der Waals interaction contributed more energy for both lipids compared to the Coulomb interaction, yet LQ-3 forged significantly more van der Waals interactions than SM-102.

Beyond the greater number of hydrogen bonds mediated by LQ-3 compared to SM-102, the occupancy of these hydrogen bonds in LQ-3-mRNA complexes is significantly elevated, as depicted by the Nightingale rose charts (Figure 4D), suggesting that these interactions in LQ-3 are more stable. A closer inspection of hydrogen bond interactions between lipid and mRNA reveals that the two hydroxyls from LQ-3 head can simultaneously form hydrogen bonds with two neighboring backbone phosphates (Figures 4E). These synergistic interactions could underline the augmented stability between LQ-3 and mRNA interactions, potentially contributing to the superior binding affinity of LQ-3 with mRNA.

### 2.6. DSPC’s Adverse Effects on LQ-3 Nanoparticle Formation and Stability

Having established LQ-3’s superior ability to complex mRNA, enabling stable particle formation without the need for DSPC, our investigation probed into the reasons behind DSPC’s adverse effects on particles, as evidenced by increased PDI and reduced mRNA delivery efficacy. We subjected both LQ-3 iPLX and LNP to stress treatments involving multiple cycles of freeze-thaw (FT). We then examined their physical-chemical properties and morphology using cryo-electron microscopy (cryo-EM). Additionally, we assessed their performance during long-term storage at −20 °C

As depicted in Figure 5A, LQ-3 iPLX demonstrated remarkable stability under both multiple freeze-thaw cycles and long-term storage at −20 °C over the observed three-month period. However, when DSPC was incorporated into the formulation, the resulting LQ-3 LNP exhibited a consistent increase in particle size and PDI, with no significant impact on encapsulation efficiency. Interestingly, for SM-102, DSPC proved to be an absolute necessity to withstand the stress treatment or long-term storage, as evidenced by the comparison between SM-102 iPLX and LNP formulations.

**Figure 5.**
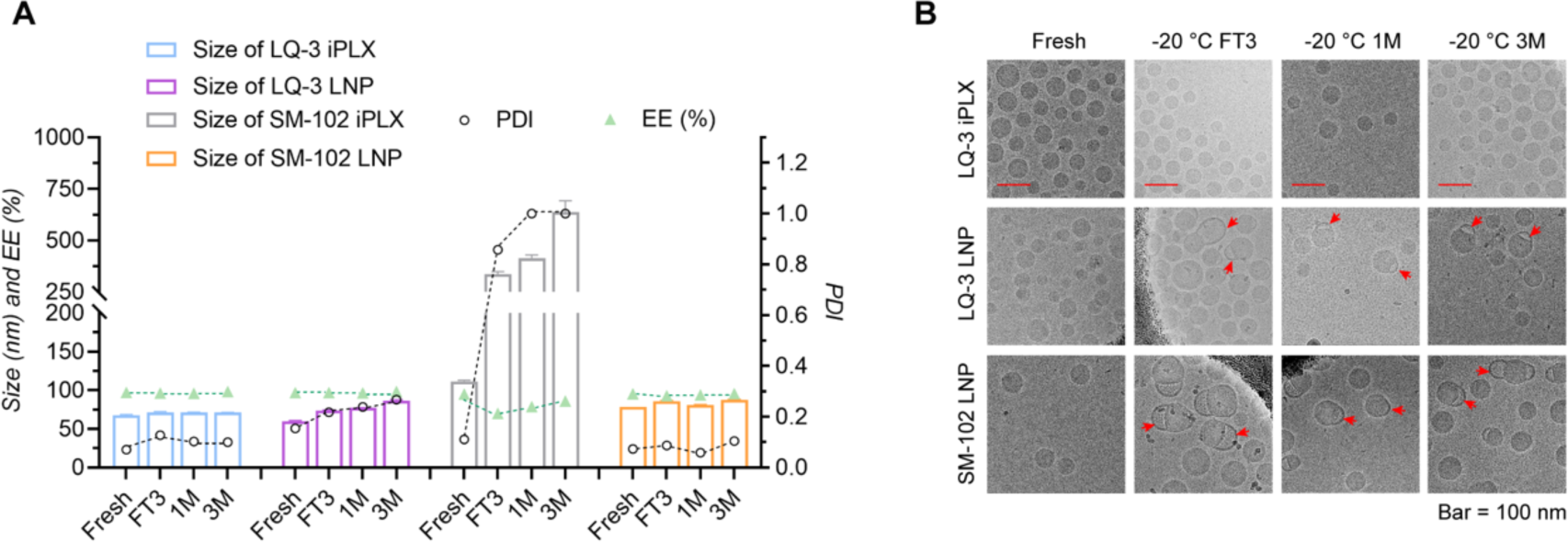
Stability assessment of LQ-3 nanoparticles. (A) Evaluation of size, PDI, and EE for iPLX and LNP following three freeze-thaw cycles (FT3) and extended storage at −20°C (n = 3, mean ± SD). (B) Cryo-EM representations of LQ-3 iPLX, LQ-3 LNP, and SM-102 LNP post sequential challenges. Red arrows indicate blebs. Scale bar measures 100 nm.

Concurring with these observations, our examination of the nanoparticles’ morphology using cryo-EM revealed that the dispersity of LQ-3 LNP was poorer than that of iPLX, with a higher number of small particles in the 20-40 nm size range within LQ-3 LNP samples (Figure 5B). Notably, no significant changes were observed in LQ-3 iPLX following either freeze-thaw treatment or long-term storage at −20 °C. However, LQ-3 LNP exhibited discernible bleb-like structures after stress treatment and one month of storage. Similar observations of bleb-like structures were also noted in SM-102 LNP following freeze-thaw treatment and long-term storage. This suggests that DSPC is the primary driver behind the formation of these blebs, in line with previous findings^27,36^.

### 2.7. Vaccine Applications of LQ-3 iPLX Platform

Up to this point, our investigations have underscored the efficiency and stability of LQ-3 iPLX as a mRNA delivery system for mRNA payloads. To harness these promising findings for practical applications, we explored the potential of iPLX as a versatile vaccine platform, specifically for developing a multivalent mRNA vaccine targeting various SARS-CoV-2 variants. We encapsulated three distinct mRNAs encoding full-length chimeric spike protein from different SARS-CoV-2 variants (B.1.1.7/B.1.351, Omicron BA.2/4/5, and XBB.1.5) into mRNA-iPLX using LQ-3 as the ionizable lipid (Figure 6A). As a reference, we also created LQ-3 empty iPLXs to serve as a negative control. In addition, we developed another mRNA vaccine that encapsulated the same mRNAs using the SM-102 LNP platform, facilitating direct comparisons. The immunogenicity of both LQ-3 iPLX and SM-102 LNP vaccines was evaluated in BALB/c mice, following the scheme outlined in Figure 6A. Mice were initially primed with intramuscular immunizations of either PBS (as a placebo), empty LQ-3 iPLX or vaccines containing varying mRNA dosages (2 μg and 5 μg) on day 0, followed by a booster injection on day 21.

**Figure 6.**
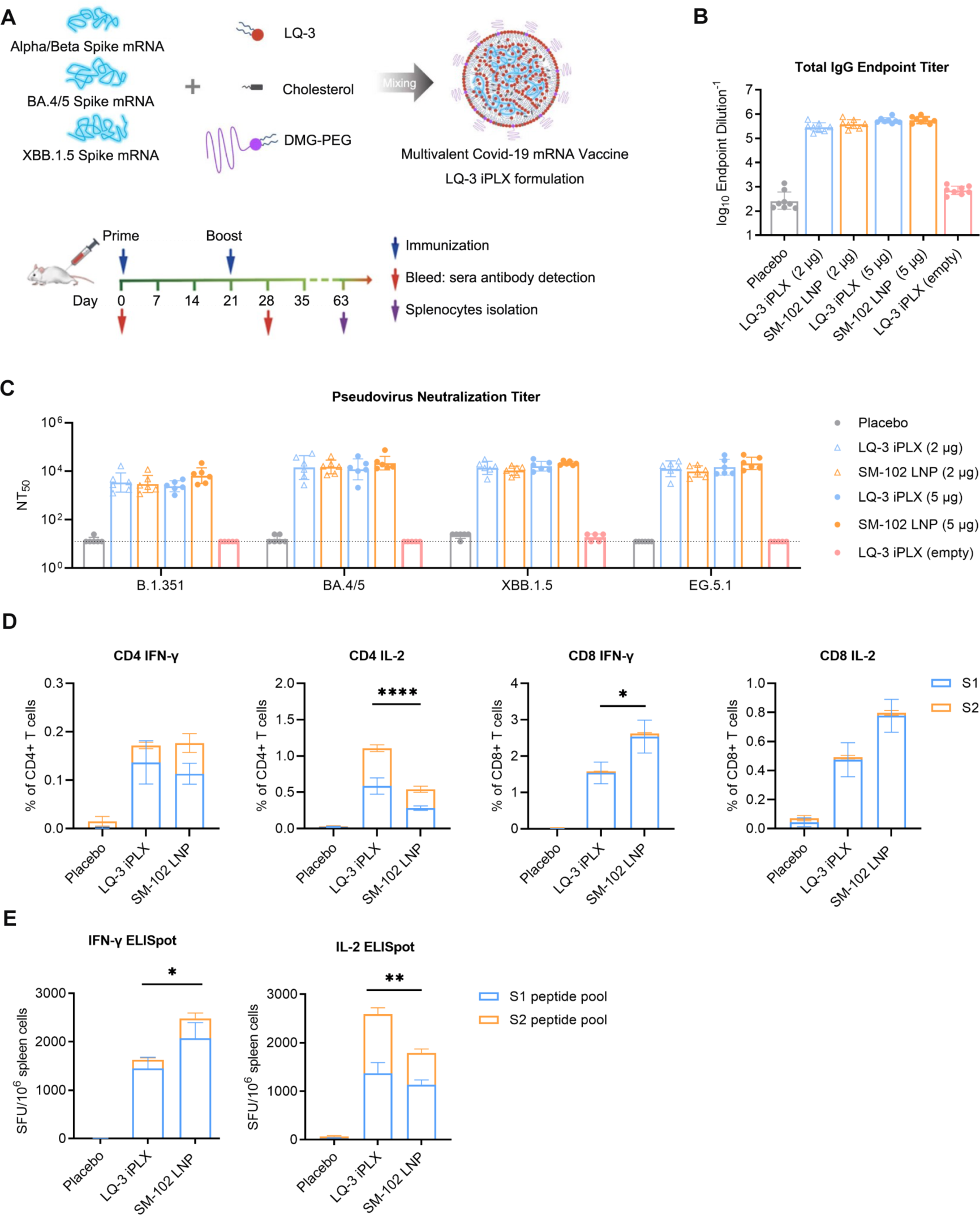
Assessment of *in vivo* immunogenicity using LQ-3 iPLX as an mRNA vaccine carrier. (A) An illustration depicting the formulation of the multivalent Covid-19 mRNA vaccine utilizing LQ-3 iPLX and the subsequent immunization strategy in mice. (B) Day 28 samples were assessed for IgG binding to the authentic SARS-CoV-2 S proteins using ELISA. Data is displayed as geometric mean ± SD. (C) Analysis of neutralizing antibody levels from day 28 sera via the lentiviral luciferase-integrated pseudovirus assay. The assay’s detection limit is marked with a black dashed line (reciprocal titer of 12.5). Data is displayed as geometric mean ± SD. (D) Analysis of CD4+ and CD8+ cells expressing IFN-γ and IL-2 was executed with intracellular cytokine staining and subsequent flow cytometry. Results are presented as mean ± SEM, with each group consisting of 6 mice. (E) Quantification of IFN-γ and IL-2 releasing cells using the ELISpot method. On day 71, splenocytes were collected stimulated using peptide mixtures from the S1 and S2 domains of the SARS-CoV-2 Spike protein. Data is presented as mean ± SEM (n = 6). Significance indicators: *p < 0.05, **p < 0.005, and ****p < 0.0001.

For subsequent evaluations, enzyme-linked immunosorbent assay (ELISA) was employed to measure S protein-specific IgG titers. As demonstrated in Figure 6B, the LQ-3 iPLX vaccine elicited a robust immune response. The S protein-specific IgG antibody titers in sera collected on day 28 from mice immunized with either 2 μg or 5 μg of LQ-3 iPLX were comparable to those in the SM-102 LNP immunized group. Furthermore, the lentiviral pseudovirus neutralization assay was employed to evaluate the levels of neutralizing antibodies (NAb) induced by the respective vaccines. Encouragingly, both the LQ-3 iPLX and SM-102 LNP vaccines induced substantial neutralization activity against the EG.5.1, XBB.1.5, Omicron BA.4/5, and B.1.351 viruses. No significant disparities were observed between these two delivery systems across both dosage groups (Figure 6C). These results compellingly suggest that our LQ-3 iPLX vaccines effectively induced a potent humoral immune response in mice.

Furthermore, we investigated the cellular immune response against the spike protein using intracellular cytokine staining (ICS) assays and ELISpot assays. The ICS results revealed that LQ-3 iPLX robustly triggered a significant increase in IFN-γ and IL-2 expression by CD8^+^ T cells compared to the placebo group (Figure 6D). The frequency of CD8+ T cells expressing IFN-γ induced by LQ-3 iPLX was slightly lower than that induced by SM-102 LNP. A similar trend was observed in the IFN-γ ELISPOT assay (Figure 6E). Additionally, mice vaccinated with LQ-3 iPLX exhibited significantly higher percentages of IL-2-specific CD4+ T cells and a greater number of IL-2-producing splenocytes compared to the SM-102 LNP-treated groups (Figure 6D). These findings firmly highlighted the capacity of the LQ-3 iPLX platform to not only elicit a strong humoral immune response in mice, but also to trigger a robust T cell immune response.

### 2.7. Safety of LQ-3 iPLX *In Vivo*

We then proceeded to assess the *in vivo* safety profile of LQ-3 iPLX encapsulating SARS-CoV-2 S protein-encoded mRNA through a single intramuscular administration in Sprague-Dawley (SD) rats and C57BL/6J mice. For the purpose of comparison, SM-102 LNP was used as a control. Rats were administered mRNA doses of 40 μg and 240 μg, respectively. Over a14-day recovery period, the immunized rats were subjected to clinical symptom observation and physiological and biochemical parameter analyses. Similarly, mice were given a mRNA dose of 20 μg, followed by a 12-day recovery period for weight monitoring and physiological and biochemical parameter analysis. Notably, no severe adverse reactions were observed in any of the animals immunized with either LQ-3 iPLX or SM-102 LNP, regardless of the administered doses. For the assessment, markers of hepatic and nephritic injury, namely alanine aminotransferase (ALT) and creatinine (Crea) levels were evaluated at 6 hours and 14 days post-administration in rats and at 12 days in mice. In comparison to the placebo group, there were no significant changes in ALT and Crea levels among rats administered either 40 μg (Figures 7A-D) or 240 μg mRNA (Figure S16). Similar findings were observed for serum ALT and Crea levels in mice 12 days after administration (Figures 7G-H).

**Figure 7.**
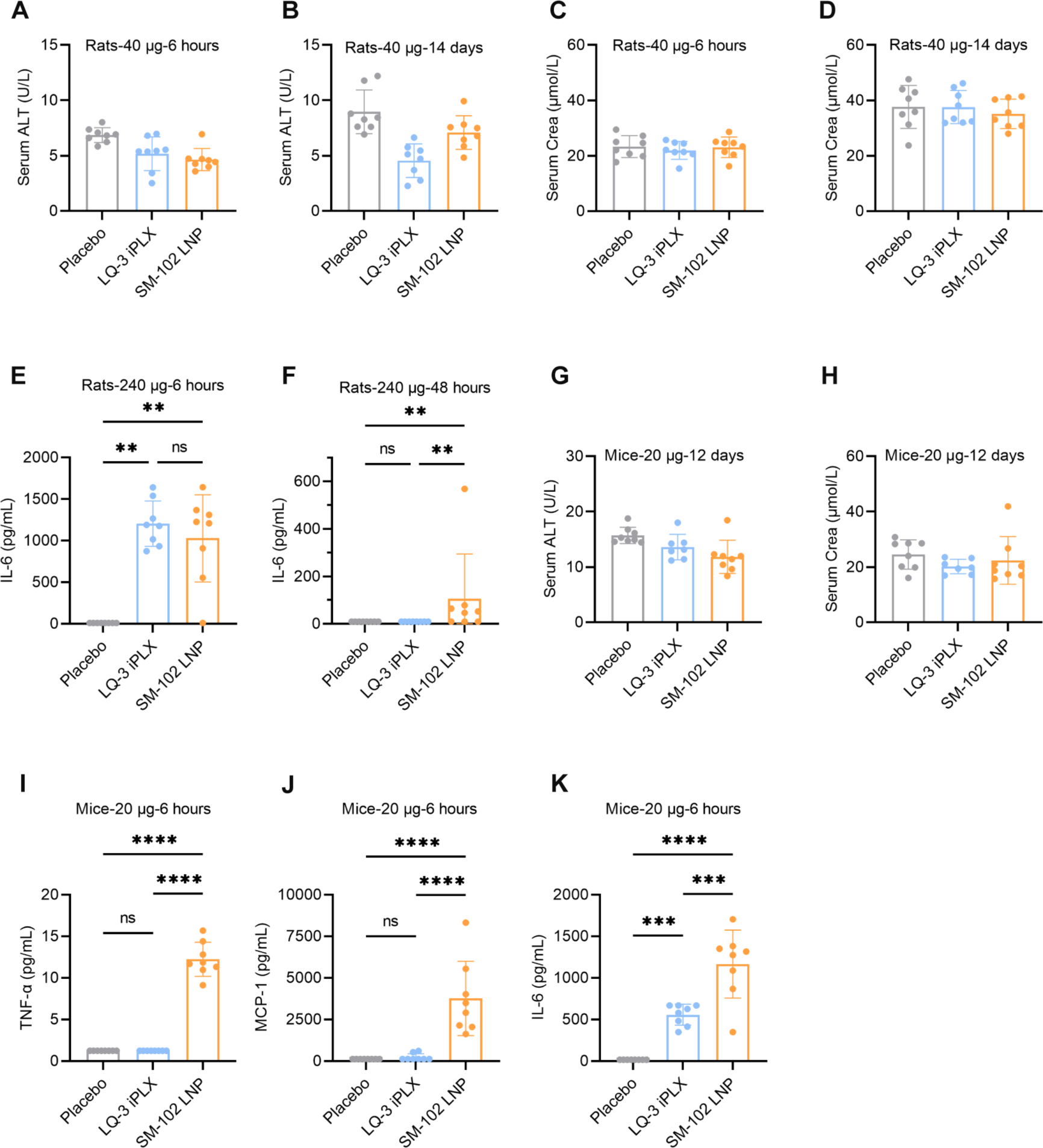
*In vivo* safety assessment of LQ-3 iPLX in rats and mice. (A-B) Serum alanine aminotransferase (ALT) levels in SD rats at 6 hours (A) and 14 days (B) post-intramuscular administration. n = 8, mean ± SD. There was no notable difference between the placebo and the nanoparticle groups at 40 μg per injection. (C-D) Serum creatinine concentrations in SD rats at 6 hours (C) and 14 days (D) following intramuscular injection. n = 8, mean ± SD. No significant variance was found between the placebo and nanoparticle cohorts at 40 μg per injection. (E-F) Serum interleukin-6 (IL-6) concentrations in SD rats at 6 hours (E) and 48 hours (F) after receiving a high dose of mRNA-LNP, as quantified by ELISA. n = 8, mean ± SD, 240 μg per injection. (G-H) ALT (G) and creatinine (H) levels in C57BL/6J mice on day 12 following intramuscular delivery. n = 8/7 (accounting for fatalities), mean ± SD. No discernible difference was identified between the placebo and nanoparticle groups at an mRNA quantity of 20 μg per injection. (I-K) Luminex assay-derived serum levels of inflammatory cytokines (TNF-α, MCP-1, and IL-6) in C57BL/6J mice 6 hours post-intramuscular injection. n = 8, mean ± SD, 20 μg per injection. Significance indicators: **p ≤ 0.01, ***p ≤ 0.001, and ****p ≤ 0.0001.

Furthermore, the innate immune response triggered by LQ-3 iPLX was examined. Six hours after administration, serum IL-6 levels in both LQ-3 iPLX and SM-102 LNP treated rats were considerably elevated compared to the placebo group (Figure 7E). Notably, while the serum IL-6 level in LQ-3 iPLX-treated rats was comparable to that in the placebo group 48 hours post-administration, the IL-6 level in the SM-102 group remained significantly higher (Figure 7F).

To gauge the levels of inflammation-related cytokines/chemokines, including TNF-α, MCP-1, IL-1β, IL-6, IL-10, and IFN-γ, a Luminex assay was conducted on mice sera. Distinct differences in cytokine activation were observed between the LQ-3 iPLX and SM-102 LNP groups at 6 hours post administration. As illustrated in Figures 7I-J, substantially elevated TNF-α and MCP-1 levels were observed in the SM-102 LNP group, in contrast to the LQ-3 iPLX group. Furthermore, neither LQ-3 iPLX nor SM-102 LNP triggered the expression of IL-1β, IL-10, or IFN-γ (Figure S17). While both LQ-3 iPLX and SM-102 LNP led to a significant increase in IL-6 expression, the IL-6 level in LQ-3 iPLX-treated mice remained lower than that in the SM-102 LNP group (Figure 7K), highlighting the considerably lower innate immune response induced by LQ-3 iPLX compared to SM-102 LNP.

Additionally, there were no notable changes in bodyweight observed in either the LQ-3 iPLX or SM-102 LNP groups when compared to the placebo group (Figure S18). However, both the LQ-3 and SM-102 groups that received a high mRNA dosage (240 μg) exhibited notably increased spleen-body weight ratios. (Figure S19). In conclusion, the preliminary toxicological studies indicated that LQ-3 iPLX system displayed favorable safety characteristics.

## 3. Discussion

The remarkable efficacy of messenger RNA (mRNA) vaccines against SARS-CoV-2 has underscored the importance of efficient mRNA delivery systems. While lipid nanoparticles (LNPs) have played a pivotal role in this success, optimizing their components remains a subject of ongoing research. The prevailing model suggests that LNP formation begins with ionizable lipids interacting with mRNA under acidic conditions, primarily driven by the ionic attraction between the positively charged ionizable lipid and the negatively charged mRNA. This interaction creates a nucleation core, which then accrues cholesterol, phospholipids, and PEG-lipid, until the surface accumulation of PEG-lipid inhibits further nanoparticle growth. Following formation, the particles are dialyzed to a neutral buffer, typically with a pH exceeding the apparent pKa of the particle. This shift in pH is believed to significantly alter the internal particle structure^25^. When the ionizable lipid loses its charge due to the elevated pH, its interaction with mRNA weakens, causing the lipid to self-aggregate and create an oil-like core phase. This reorganization propels mRNA towards the periphery of the LNP near the shell, as revealed by cryo-EM imaging (Figure 3B, See also references^25,35^). It is assumed that phospholipids, predominantly situated on the shell, are key in keeping mRNA encapsulated. Intriguingly, LNP particles tend to morph into bleb-like structures when phospholipid DSPC is employed, as DSPC has a strong propensity to form a lipid bilayer structure that would segregate into blebs under stress or over extended storage (Figure 5B), which is usually followed by the release of mRNA into the bleb space^35^. When the ionizable lipid reacquires its charge via pH reduction, mRNA is reabsorbed into the lipid core. This dynamic underscores the crucial role of ionic interaction in maintaining LNP stability.

In our study, we unveiled a novel series of ionizable lipids characterized by double ethanolamine head groups with two biodegradable branched tails, exemplified by the leading compound LQ-3. The overall mRNA delivery efficiency of LQ-3 LNP was comparable to that of lipids with a single ethanolamine head, like SM-102. However, a unique advantage of LQ-3 became apparent when we attempted to optimize its formulation -- including phospholipids like DSPC or DOPE negatively impacted delivery and compromised particle stability in LQ-3 formulations, a stark contrast to SM-102. We speculate that these disparities likely arise from the head configuration of LQ-3, implying that the double ethanolamine head might possess a higher affinity for mRNA. This might be due to the two positive charges carried by the LQ-3 head compared to one by SM-102, or LQ-3 might be more adept at forming nonionic interactions, such as hydrogen bonds with mRNA. To differentiate these interactions, we mixed LQ-3 with mRNA under acidic or neutral conditions and discovered that LQ-3 rendered over 95% mRNA inaccessible to RiboGreen dye, irrespective of the pH, indicating a major role of nonionic interactions (Figure 3D). This was further substantiated by our molecular dynamics (MD) simulation, revealing that the hydroxyl groups in LQ-3’s head establish an increased number of hydrogen bonds with mRNA, often synergistically (Figure 4).

These dominant nonionic interactions entail two significant implications. Firstly, the dependence on the charges of ionizable lipids during the initial nanoparticle formation phase is lessened, as formation can now proceed in a neutral buffer without compromising encapsulation efficiency. This is advantageous during LNP manufacturing, as the process can be executed in the same buffer, avoiding pH adjustments which frequently disrupt particle structure leading to aggregation. Secondly, the “helper” phospholipids either lose their importance or negatively affect particle stability and delivery efficiency. As detailed earlier, phospholipids like DSPC are essential for SM-102 and possibly other ionizable lipids to form stable particles, as they could mitigate the adverse impact triggered by the phase separation of ionizable lipid under neutral conditions. However, in the case of ionizable lipid like LQ-3, where nonionic interactions predominate, the RNA:lipid core remains relatively stable even when ionizable lipids lose charges, as inferred from cryo-EM images (Figure 3B). Hence, DSPC is not imperative for stability. This is reminiscent of conventional lipoplex formulation, where a permanently charged cationic lipid is used to complex with DNA or oligodeoxynucleotides, and stable particles can be formed by supplying the auxiliary lipid cholesterol^34^.

Building on these insights, we developed a new iPLX formulation encompassing three components: ionizable lipid LQ-3, cholesterol, and PEG-lipid. In practice, we observed that LQ-3 iPLX generally achieved a higher mRNA encapsulation efficiency of around 98%, while SM-102 LNP usually yielded around 92% efficiency. Additionally, the exclusion of DSPC also eliminated the formation of the bleb-like structure (Figure 5B).

Comparing mRNA expression, the peak expression of mRNA-LQ-3 iPLX at the 4-hour mark slightly lagged behind mRNA-SM-102 LNP, but it waned at a slower rate from 4 to 24 hours. The rationale for this could be a less productive endosome escape of mRNA, as DSPC has been implicated in promoting endosome escape, or stronger binding of ionizable lipids to mRNA post-endosome escape, interfering with the ribosome from engaging mRNA for translation initiation. Further studies are needed to address these questions and may help to guide further optimization of the ionizable lipid.

We also explored the potential of LQ-3 iPLX as a vaccine platform. When compared with SM-102 LNP, we noticed that LQ-3 iPLX elicits similar levels of humoral response, however, T-cell immune response is slightly weaker. Further analysis revealed that LQ-3 iPLX induced significantly lower pro-inflammatory cytokines such as IL-6 and TNF-α, which could account for the weaker cell immune response (Figure 7). Whether this is due to the absence of DSPC or lesser adjuvant activity of LQ-3 compared to SM-102 is yet to be determined. Notably, owing to the two amine groups in LQ-3, a reduced quantity of LQ-3 was required to deliver the same amount of RNA compared to SM-102 with a fixed N:P ratio. Nevertheless, LQ-3 iPLX holds promise for a more controllable immune response, potentially reducing the prevalent adverse effects of current mRNA vaccines on the market.

In conclusion, our research has successfully uncovered a novel series of ionizable lipids that facilitate a three-component formulation, paving the way for enhanced stability and efficacy in mRNA delivery. This advancement also marks a significant stride towards lipid optimization. By capitalizing on the non-electrostatic interactions between ionizable lipids and mRNA, we can now tailor the formulation for more simplified and safer delivery systems.

## 3. Experimental Section

### mRNA Synthesis

The mRNA encoding spike protein (Spike mRNA) and mRNA encoding firefly luciferase protein (Fluc mRNA) were synthesized in vitro by T7 RNA polymerase-mediated transcription from a linearized DNA template. Cap1 was utilized to increase mRNA translation efficiency.

### Lipid synthesis

A series of compounds were synthesized and their structures were confirmed following the procedures outlined in the Supplemental Materials (Figures S1-S14). SM-102 was synthesized following the previously reported procedure^14^. DSPC and Cholesterol were purchased from Nippon Fine Chemical Co., Ltd. DMG-PEG2k was obtained from Guobang Pharma Ltd.

### Preparation of mRNA-encapsulated lipid nanoparticles

mRNA-loaded LNPs were prepared by formulating a total lipid concentration of 12.5 mM, consisting of the synthetized ionizable lipids/DSPC/Cholesterol/DMG-PEG2k at specific mole ratios (refer to Tables S1-S3). The individual lipids were dissolved in ethanol and carefully mixed to attain the desired molar ratios. The Fluc mRNA or Spike mRNA was dissolved in a 25 mM sodium acetate buffer with a pH of 5.0 (or a 50 mM citric acid buffer with a pH of 4.0). The two phases were rapidly mixed through microfluidic device at a volume ratio of 3:1 (mRNA: lipid) with a total flow rate of 12 mL/min. The resulting formulations were dialyzed against 10 mM PBS (pH 7.2∼7.4) or 20 mM Tris-Acetate (pH ∼7.5) using Slide-A-Lyzer dialysis cassettes. Following dialysis, the solutions were concentrated using Amicon ultra-centrifugal filters and filtered through a 0.22 μm filter.

### LNP characterization

The formulations were evaluated for particle size, RNA encapsulation and pKa. The average particle diameter and polydispersity index (PDI) were determined using Zetasizer Nano ZS (Malvern Instruments Ltd., UK). The encapsulation efficiency (EE) of mRNA was measured using the Quant-it Ribogreen Assay Kit (Thermo Fisher Scientific, USA). The *p*Ka of LNP was determined using the TNS binding assay^14^.

### mRNA binding affinity

LQ-3 and SM-102 was dissolved in ethanol at a concentration of 6.25 mM respectively. A certain concentration of mRNA was dissolved in a 25 mM sodium acetate buffer with a pH of 5. The two phases were rapidly mixed at a volume ratio of 3:1 (mRNA: lipid). Maintain the original solution or adjust the system pH to near neutral. The binding affinity of mRNA at two pH values was measured using the Quant-it Ribogreen Assay Kit (Thermo Fisher Scientific, USA).

### 1H-NMR

The surface lipid distribution of LNP was characterized using an ultra-high field (800 MHz) nuclear magnetic resonance spectrometer (NMR). The LNPs were dialyzed into 0.2 × PBS to remove sucrose. By combining the chemical structure of each lipid and utilizing one-dimensional and two-dimensional NMR spectra, the spectral peak of LNP was identified through chemical shift analysis (MestReNova 14.2 software).

### Cryo-EM

Cryo-EM specimens were prepared at the original concentration of LNP on the ANTcryoTM Au300 R1.2/1.3 grids (Nanodim). The grids were glow-discharged using an easyGlow device (PELCO) with the parameters set at 15 mA and 100 seconds. A Vitrobot plunge freezer (Vitrobot Mark IV, Thermo Fisher Scientific) was used for preparing cryo grids. A droplet of 3 ul was applied to the freshly glow-discharged grid and the grid was blotted (bolt time 4 s, bolt force 0, wait time 30 s) and plunge-frozen in liquid ethane then stored in liquid nitrogen until usage. Autogrids were assembled under liquid nitrogen and then loaded into the electron microscope for further examination. Data collection was carried out on a Glacios transmission electron microscope (Thermo Fisher Scientific) operated at 200 kV, equipped with a Ceta-D camera. Images were taken using the EPU software (Thermo Fisher Scientific).

### Freeze-thaw challenges

LNPs were prepared and dispersed in tris-ac buffer supplemented with a sucrose protectant, resulting in a final mRNA concentration of 200 μg/mL. In each freeze-thaw cycle, the freeze-thaw procedure entailed subjecting the LNPs to a freezing temperature of −20°C for a minimum of 20 hours, followed by thawing at 4°C for approximately 2 hours. The particle size, PDI, mRNA encapsulation efficiency was determined, and morphology of the LNPs were assessed using a Cryo-EM electron microscope.

### Molecular dynamics

MD trajectories were acquired using Gromacs2021 package, GPU-accelerated, employing the AMBER14SB and GAFF all-atom forcefields^37,38^. Each simulation box, maintained at a 1:1 base to lipid ratio, housed a 28-nucleotide mRNA, 28 ionizable lipids either SM-102 or LQ-3, and was solvated with explicit TIP3P water molecules within a vacuum box placed 5 nm away from the border. The systems were initialized with bond length, angle, and dihedral values for the lipids generated using Sobtop, and the RESP charges computed via Gaussian and Multiwfn^39^. Following energy minimization using a steepest descent approach, a pre-equilibration phase was conducted through a 500 ps isothermal-isobaric simulation (NPT). The subsequent 100 ns production MD simulation was carried out, with the operational parameters set to maintain isotropic coupling, a compressibility of 4.5 × 10^−5^, and a coupling constant of 2.0 ps, in alignment with the Parrinello−Rahman algorithm. The temperature was held steady at 298.15K, with a coupling constant of 0.2 ps as per the v-rescale algorithm. Short-range interactions were set with a 1.0 nm cutoff, while the Particle Mesh Ewald algorithm was employed for long-range electrostatic interactions. Trajectory snapshots were recorded every 10 ps, documenting dynamic integrals at 2 fs intervals. Analysis and visualization of the motion trajectories, hydrogen bond characteristics, and conformational changes were facilitated using Visual Molecular Dynamics^40^ and Qtgrace tools.

### Animals

Female BALB/c mice weighing 18-22 g were obtained from Sibeifu (Beijing) Biotechnology Co., Ltd. C57BL/6J mice weighing 19-24 g were obtained from Shanghai Lingchang Biotechnology Co., Ltd. Sprague-Dawley rats weighing 225-250 g were purchased from Shanghai Jihui Experimental Animal Breeding Co., Ltd. All animals were given unrestricted access to water and food. Animal experiments were conducted in accordance with the Principles of Experimental Animal Ethics.

### *In vivo* screening

In vivo live animal imaging for luciferase expression following intravenous administration of LNPs encapsulating FLuc mRNA at a dose of 5 μg mRNA/ injection. The imaging procedure was performed at 4-hour, 24-hour, and 48-hour post-injection. Mice received an intraperitoneal injection of 100 μL D-Luciferin (30 mg/mL) 8 min before imaging. Bioluminescence imaging was performed using an IVIS Spectrum imaging system (PerkinElmer). PK curve and area under the curve (AUC) was analyzed by GraphPad Prism 9 software.

### Animal vaccination and serum collection

For mouse vaccination, groups of 6- to 8-week-old female BALB/c were intramuscularly immunized with LNP vaccine candidates or a placebo in 50 μL, and 3 weeks later, a second dose was administered to boost the immune responses.

### *In vivo* safety evaluation of LNPs in rat and mouse

The tolerability *in vivo* of LQ-3 was evaluated in Sprague-Dawley rats and C57BL/6j mice. LQ-3 iPLX encapsulating Spike mRNA was administered to mice and rats via intramuscular injection. Tris-Ac buffer placebo was used as the negative control, and SM-102 LNP encapsulating the Spike mRNA was utilized as the positive control. The dose of mRNA-LNP given to mice was 20 μg mRNA/injection, with 8 female mice in each group. The day of administration was defined as day 0. The serum samples were collected at 6 hours and day 12 after administration to detect cytokines and serum markers of hepatic and nephritic injury, such as ALT and creatinine. The body weight of each group of mice was recorded on day 0, 1, 2, 4, 6, 8, 11, and 12. SD rats received doses of 40 μg mRNA/injection and 240 μg mRNA/ injection. Each group consisted of 4 female and 4 male rats. Serum samples were collected at 6 hours, 48 hours, and 14 days after administration to measure cytokines, ALT, and creatinine. The rats were monitored for behavior and appearance during a 14-day recovery period after administration. A gross dissection was performed at day 14 to observe the appearance and record the weight of heart, liver, spleen, lung and kidney.

### ELISA for SARS-CoV-2 S-specific IgG

SARS-CoV-2 S-specific antibody responses in immunized sera were determined by enzyme-linked immunosorbent assay (ELISA) assay, as previously described. Briefly, 96-well plates were coated with 50 μL of coating buffer containing 100 ng/well recombinant SARS-CoV-2 spike antigens (Sino Biological) at 4 °C overnight. Plates were blocked with 2% bovine serum albumin solution in PBST at room temperature for 1 hour. Immunized mice sera were diluted 100-fold as the initial concentration, and then a 5-fold serial dilution of a total of 11 gradients in PBS buffer. PBST washed plates were incubated with serially diluted sera at room temperature for 2 hours. For determination of S-specific antibody response, plates were incubated with goat anti-mouse IgG HRP (Proteintech Cat: SA00001-1) at 37 °C for 1 hour and then substrate tetramethylbenzidine (TMB) solution (Invitrogen) was used to develop. The color reaction was quenched with 1N sulfuric acid for about 10 minutes, and the optical density was measured at a wavelength 450 nm by VARIOSKAN LUX (Thermo Fisher).

### Pseudovirus-based neutralization assay

Serum samples collected from immunized animals were serially diluted with the cell culture medium. The diluted serums were mixed with a pseudovirus suspension in 96-well plates at a ratio of 1:1, followed by 1 hour incubation at RT. Opti-HEK293/ACE2 cells were then added to the serum-virus mixture, and the plates were incubated at 37 °C in a 5% CO_2_ incubator. 48 hours later, the luciferase activity, reflecting the degree of SARS-CoV-2 pseudovirus transfection, was measured using the Luciferase Assay kit. The NT_50_ was defined as the fold-dilution, which emitted an exceeding 50% inhibition of pseudovirus infection compared to the control group.

### Intracellular cytokine staining assay

For the intracellular cytokine staining, mouse spleen cells were stimulated with 2 μg/ml S1、S2 peptide pools spanning SARS-CoV-2 spike S1 and S2 respectively (15mers, overlapping by 11aa, GenScript) or equimolar amount of DMSO (negative control) in the presence of anti-mouse CD28 antibody (BD Bioscience) for 1 hour. GolgiStop™ protein transport inhibitor was added into the culture and further incubated for 5 hours. After stimulation, cells were washed and stained with Fixable Viability Stain 510 (BD Bioscience). Cells were then blocked with anti-mouse CD16/32 and labeled with cell surface antibody cocktail including anti-mouse CD3-FITC、anti-mouse CD4-APC and anti-mouse CD8-Percp-cy5.5 (BD Bioscience) for 30min at 4°C. Following surface staining, cells were incubated with fixed/permeabilized solution (BD Bioscience) for 20min at 4°C and stained with intracellular antibodies including anti-mouse IFN-γ-Pe-Cy7 and IL-2-BV605 (BD Bioscience) for 30min at 4°C. Cells were washed and resuspended in PBS buffer and acquired on a CYTEK Aurora/NL.

### IFN-γ and IL-2 ELISpot assays

The number of spleen cells secreting IFN-γ and IL-2 was detected by enzyme-linked immunospot (ELISpot) assays. In brief, single cell suspension from the mouse spleen was prepared and stimulated with 2 μg/ml spike S1、S2 peptide pool or equimolar amount of DMSO(negative control) for 22 h. IFN-γ and IL-2-secreting cells were detected using ELISpot kits (DAKEWE and MABTECH) according to the manufacture protocol and spots were counted using an ELISpot Analyzer AT-SPOT(SINSAGE).

### Luminex assay for mouse serum cytokines/chemokine detection

Mouse serum cytokine/chemokine levels were measured using the Mouse Premixed Multi-Analyte Kit (R&D Systems, USA) at 6 hours after administration. The cytokines analyzed included TNF-α, MCP-1, IL-1β, IL-6, IL-10, and IFN-γ. The specific procedures were conducted according to the manufacturer’s recommendations. Serum samples were diluted 2-fold for measurement. Plates were read on a Luminex®200™ instrument. The lower bound of per sample per cytokine/chemokine was set as 50 beads. Values below the limit of quantification (LOQ) were assigned as half of the LOQ.

### ELISA for rat serum IL-6 detection

To detect Rat IL-6 levels, the serum samples collected at 6 hours and 48 hours after administration were diluted 2-fold for ELISA analysis using the LEGEND MAX™ Rat IL-6 ELISA Kit with pre-coated plate (Biolegend, USA) according to the manufacturer’s recommendations with a little modification as described below. In the final color development step, the reaction was stopped after adding Substrate Solution F for 5 minutes instead of 10 minutes, to avoid the results outside the instrument measuring range. The plates were read on a microplate reader, varioskan lux (therm23hermosntific), at the absorbance of 450 nm. Values below the LOQ were assigned as half of the LOQ.

### Measurements of serum ALT and creatinine levels

ALT activity and creatinine (Crea) concentration in the serum of SD rats and C57BL/6J mice collected at 6 hours and 14 or 12 days after administration were determined using the ALT assay kit and Creatinine assay kit, respectively, following the instructions provided by the reagent kits (Nanjing Jiancheng Bioengineering Institute, China).

## Author Contributions

J.Lin developed the lipid molecules; J.Lu and T.J. designed the experiments; T.J. performed physicochemical characterization of nanoparticles; L.C. and J.G. synthesized the lipid compounds; X.L. designed and conducted the safety assessments of the delivery system; S.T. assisted in the design of immunogenicity tests; H.G. supported cellular immunity detection; R.Y. aided nanoparticle preparation and characterization; Y.Z., J.Z., and D.O. conducted molecular dynamics simulations; H.Yao supported NMR studies; T.J., J.Lu, and J.Lin analyzed the data; T.J. drafted the manuscript; J.Lin, J. Lu, and M.Q. revised the manuscript; J.Lin, J.Lu, M.Q., and H.Yu conceived and supervised the project. The final manuscript was edited and approved by all authors.

## Data and materials availability

All data associated with this study are present in the paper and/or the Supplementary Materials.

## References

1 Creech, C. B., Walker, S. C. & Samuels, R. J. SARS-CoV-2 vaccines. JAMA 325, 1318–1320 (2021). 10.1001/jama.2021.3199

2 Chaudhary, N., Weissman, D. & Whitehead, K. A. mRNA vaccines for infectious diseases: principles, delivery and clinical translation. Nat. Rev. Drug Discov. 20, 817–838 (2021). 10.1038/s41573-021-00283-5

3 Huang, X. G. et al. The landscape of mRNA nanomedicine. Nat. Med. 28, 2273–2287 (2022). 10.1038/s41591-022-02061-1

4 Pardi, N., Hogan, M. J. & Weissman, D. Recent advances in mRNA vaccine technology. Curr. Opin. Immunol. 65, 14–20 (2020). 10.1016/j.coi.2020.01.008

5 Rohner, E., Yang, R., Foo, K. S., Goedel, A. & Chien, K. R. Unlocking the promise of mRNA therapeutics. Nat. Biotechnol. 40, 1586–1600 (2022). 10.1038/s41587-022-01491-z

6 Hou, X. C., Zaks, T., Langer, R. & Dong, Y. Z. Lipid nanoparticles for mRNA delivery. Nat. Rev. Mater. 6, 1078–1094 (2021). 10.1038/s41578-021-00358-0

7 Akinc, A. et al. The Onpattro story and the clinical translation of nanomedicines containing nucleic acid-based drugs. Nat. Nanotechnol. 14, 1084–1087 (2019). 10.1038/s41565-019-0591-y

8 Hald Albertsen, C., et al. The role of lipid components in lipid nanoparticles for vaccines and gene therapy. Adv Drug Deliv Rev 188, 114416 (2022). 10.1016/j.addr.2022.114416

9 Han, X. X. et al. An ionizable lipid toolbox for RNA delivery. Nat. Commun. 12, 7233 (2021). 10.1038/s41467-021-27493-0

10 Zhang, Y. B., Sun, C. Z., Wang, C., Jankovic, K. E. & Dong, Y. Z. Lipids and lipid derivatives for RNA delivery. Chem. Rev. 121, 12181–12277 (2021). 10.1021/acs.chemrev.1c00244

11 Qiu, M., Li, Y. M., Bloomer, H. & Xu, Q. B. Developing biodegradable lipid nanoparticles for intracellular mRNA delivery and genome editing. Accounts Chem. Res. 54, 4001–4011 (2021). 10.1021/acs.accounts.1c00500

12 Qiu, M. et al. Lipid nanoparticle-mediated codelivery of Cas9 mRNA and single-guide RNA achieves liver-specific in vivo genome editing of Angptl3. Proc Natl Acad Sci U S A 118, e2020401118 (2021). 10.1073/pnas.2020401118

13 Qiu, M. et al. Lung-selective mRNA delivery of synthetic lipid nanoparticles for the treatment of pulmonary lymphangioleiomyomatosis. Proc Natl Acad Sci U S A 119, e2116271119 (2022). 10.1073/pnas.2116271119

14 Sabnis, S. et al. A novel amino lipid series for mRNA delivery: improved endosomal escape and sustained pharmacology and safety in non-human primates. Mol Ther 26, 1509–1519 (2018). 10.1016/j.ymthe.2018.03.010

15 Lam, K. et al. Unsaturated, trialkyl ionizable lipids are versatile lipid-nanoparticle components for therapeutic and vaccine applications. Adv Mater 35, e2209624 (2023). 10.1002/adma.202209624

16 Hashiba, K. et al. Branching ionizable lipids can enhance the stability, fusogenicity, and functional delivery of mRNA. Small Sci. 3, 2200071 (2023). 10.1002/smsc.202200071

17 Miao, L. et al. Delivery of mRNA vaccines with heterocyclic lipids increases anti-tumor efficacy by STING-mediated immune cell activation. Nat. Biotechnol. 37, 1174–1182 (2019). 10.1038/s41587-019-0247-3

18 Cornebise, M. et al. Discovery of a novel amino lipid that improves lipid nanoparticle performance through specific interactions with mRNA. Advanced Functional Materials 32, 2106727 (2021). 10.1002/adfm.202106727

19 Belliveau, N. M. et al. Microfluidic synthesis of highly potent limit-size lipid nanoparticles for in vivo delivery of siRNA. Mol. Ther.-Nucl. Acids 1, e37 (2012). 10.1038/mtna.2012.28

20 Suk, J. S., Xu, Q. G., Kim, N., Hanes, J. & Ensign, L. M. PEGylation as a strategy for improving nanoparticle-based drug and gene delivery. Advanced Drug Delivery Reviews 99, 28–51 (2016). 10.1016/j.addr.2015.09.012

21 Cheng, X. & Lee, R. J. The role of helper lipids in lipid nanoparticles (LNPs) designed for oligonucleotide delivery. Adv. Drug Deliv. Rev. 99, 129–137 (2016). 10.1016/j.addr.2016.01.022

22 Wheeler, J. J. et al. Stabilized plasmid-lipid particles: construction and characterization. Gene Ther. 6, 271–281 (1999). 10.1038/sj.gt.3300821

23 Chander, N., Basha, G., Yan Cheng, M. H., Witzigmann, D. & Cullis, P. R. Lipid nanoparticle mRNA systems containing high levels of sphingomyelin engender enhanced protein expression in hepatic and extra-hepatic tissues. Molecular Therapy - Methods & Clinical Development 30, 235–245 (2023). 10.1016/j.omtm.2023.06.005

24 Cheng, M. H. Y. et al. Induction of bleb structures in lipid nanoparticle formulations of mRNA leads to improved transfection potency. Adv. Mater. 35, 2303370 (2023). 10.1002/adma.202303370

25 Kulkarni, J. A. et al. On the formation and morphology of lipid nanoparticles containing ionizable cationic lipids and siRNA. Acs Nano 12, 4787–4795 (2018). 10.1021/acsnano.8b01516

26 Li, J. et al. A review on phospholipids and their main applications in drug delivery systems. Asian J. Pharm. Sci. 10, 81–98 (2015). 10.1016/j.ajps.2014.09.004

27 Leung, A. K. K., Tam, Y. Y. C., Chen, S., Hafez, I. M. & Cullis, P. R. Microfluidic mixing: A general method for encapsulating macromolecules in lipid nanoparticle systems. J. Phys. Chem. B 119, 8698–8706 (2015). 10.1021/acs.jpcb.5b02891

28 Schoenmaker, L. et al. mRNA-lipid nanoparticle COVID-19 vaccines: Structure and stability. Int. J. Pharm. 601, 120586 (2021). 10.1016/j.ijpharm.2021.120586

29 Eygeris, Y., Gupta, M., Kim, J. & Sahay, G. Chemistry of lipid nanoparticles for RNA delivery. Accounts Chem. Res. 55, 2–12 (2022). 10.1021/acs.accounts.1c00544

30 Fenton, O. S. et al. Bioinspired alkenyl amino alcohol ionizable lipid materials for highly potent in vivo mRNA delivery. Adv. Mater. 28, 2939–2943 (2016). 10.1002/adma.201505822

31 Dilliard, S. A., Cheng, Q. & Siegwart, D. J. On the mechanism of tissue-specific mRNA delivery by selective organ targeting nanoparticles. Proc. Natl. Acad. Sci. U. S. A. 118, e2109256118 (2021). 10.1073/pnas.2109256118

32 Jayaraman, M. et al. Maximizing the potency of siRNA lipid nanoparticles for hepatic gene silencing in vivo. Angew. Chem.-Int. Edit. 51, 8529–8533 (2012). 10.1002/anie.201203263

33 Kulkarni, J. A., Witzigmann, D., Leung, J., Tam, Y. Y. C. & Cullis, P. R. On the role of helper lipids in lipid nanoparticle formulations of siRNA. Nanoscale 11, 21733–21739 (2019). 10.1039/c9nr09347h

34 Templeton, N. S. et al. Improved DNA: liposome complexes for increased systemic delivery and gene expression. Nat Biotechnol 15, 647–652 (1997). 10.1038/nbt0797-647

35 Brader, M. L. et al. Encapsulation state of messenger RNA inside lipid nanoparticles. Biophys. J. 120, 2766–2770 (2021). 10.1016/j.bpj.2021.03.012

36 Szebeni, J. et al. Insights into the structure of comirnaty covid-19vaccine: A theory on soft, partially bilayer-covered nanoparticles with hydrogen bond-stabilized mRNA-lipid complexes. Acs Nano 17, 13147–13157 (2023). 10.1021/acsnano.2c11904

37 Schmid, N. et al. Definition and testing of the GROMOS force-field versions 54A7 and 54B7. Eur Biophys J 40, 843–856 (2011). 10.1007/s00249-011-0700-9

38 Van Der Spoel, D. et al. GROMACS: fast, flexible, and free. J Comput Chem 26, 1701–1718 (2005). 10.1002/jcc.20291

39 Lu, T. & Chen, F. Multiwfn: a multifunctional wavefunction analyzer. J Comput Chem 33, 580–592 (2012). 10.1002/jcc.22885

40 Humphrey, W., Dalke, A. & Schulten, K. VMD: visual molecular dynamics. J Mol Graph 14, 33–38, 27-38 (1996). 10.1016/0263-7855(96)00018-5

