## Supporting Information for "Dual Ethanolamine Head Groups in Ionizable Lipids Facilitate Phospholipid-free Stable Nanoparticle Formulation for Augmented and Safer mRNA Delivery"

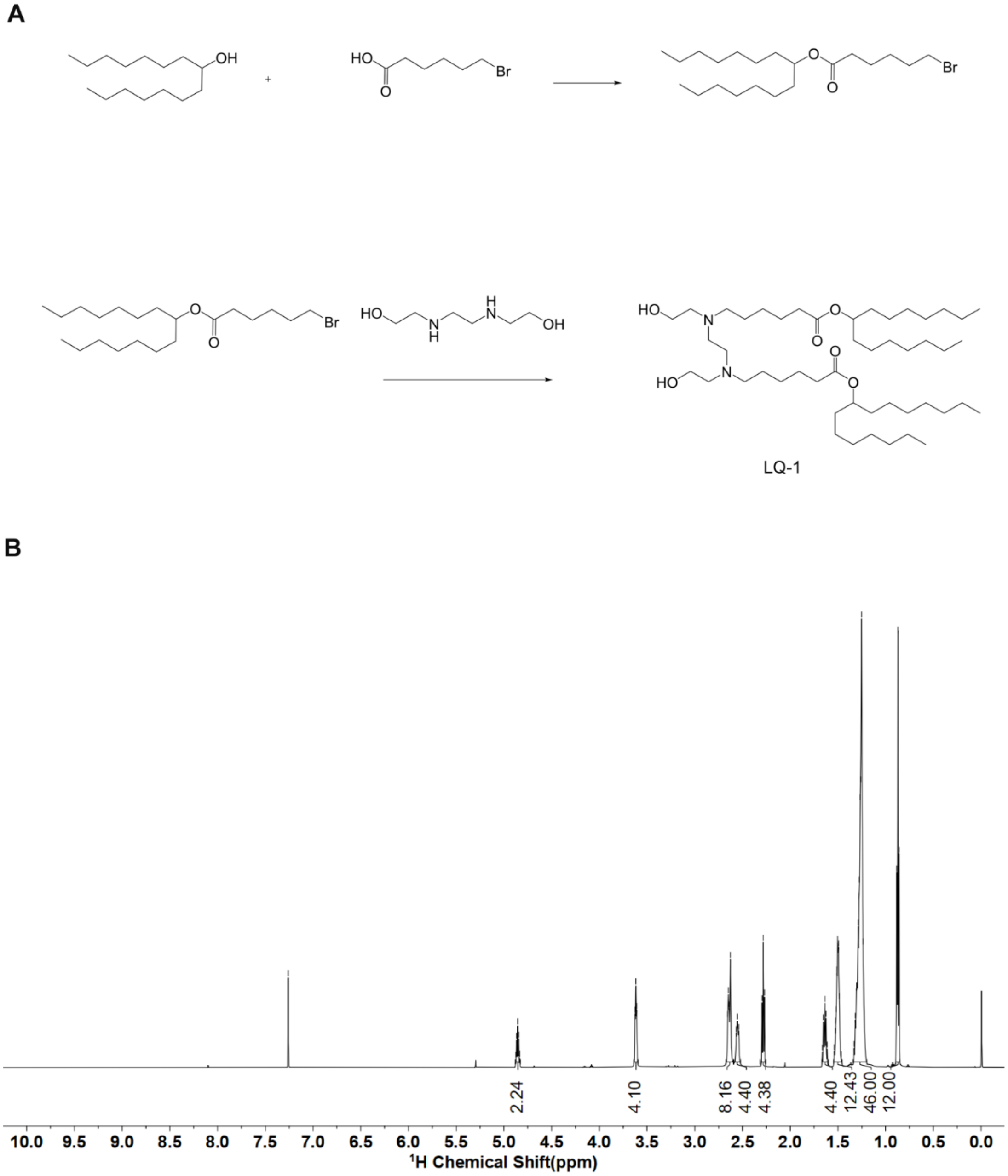


**Figure S1. The synthetic route (A) and $^{1}$H NMR (B) of LQ-1.** $^{1}$H NMR (600 MHz, $CDCl_3$) δ： 4.86 (p, *J* = 6.2 Hz, 2H), 3.62 (t, *J* = 4.9 Hz, 4H), 2.67 – 2.59 (m, 8H), 2.55 (t, *J* = 8.0 Hz, 4H), 2.28 (t, *J* = 7.5 Hz, 4H), 1.64 (p, *J* = 7.5 Hz, 4H), 1.50 (tt, *J* = 8.5, 4.5 Hz, 12H), 1.34 – 1.20 (m, 46H), 0.87 (t, *J* = 7.0 Hz, 12H).

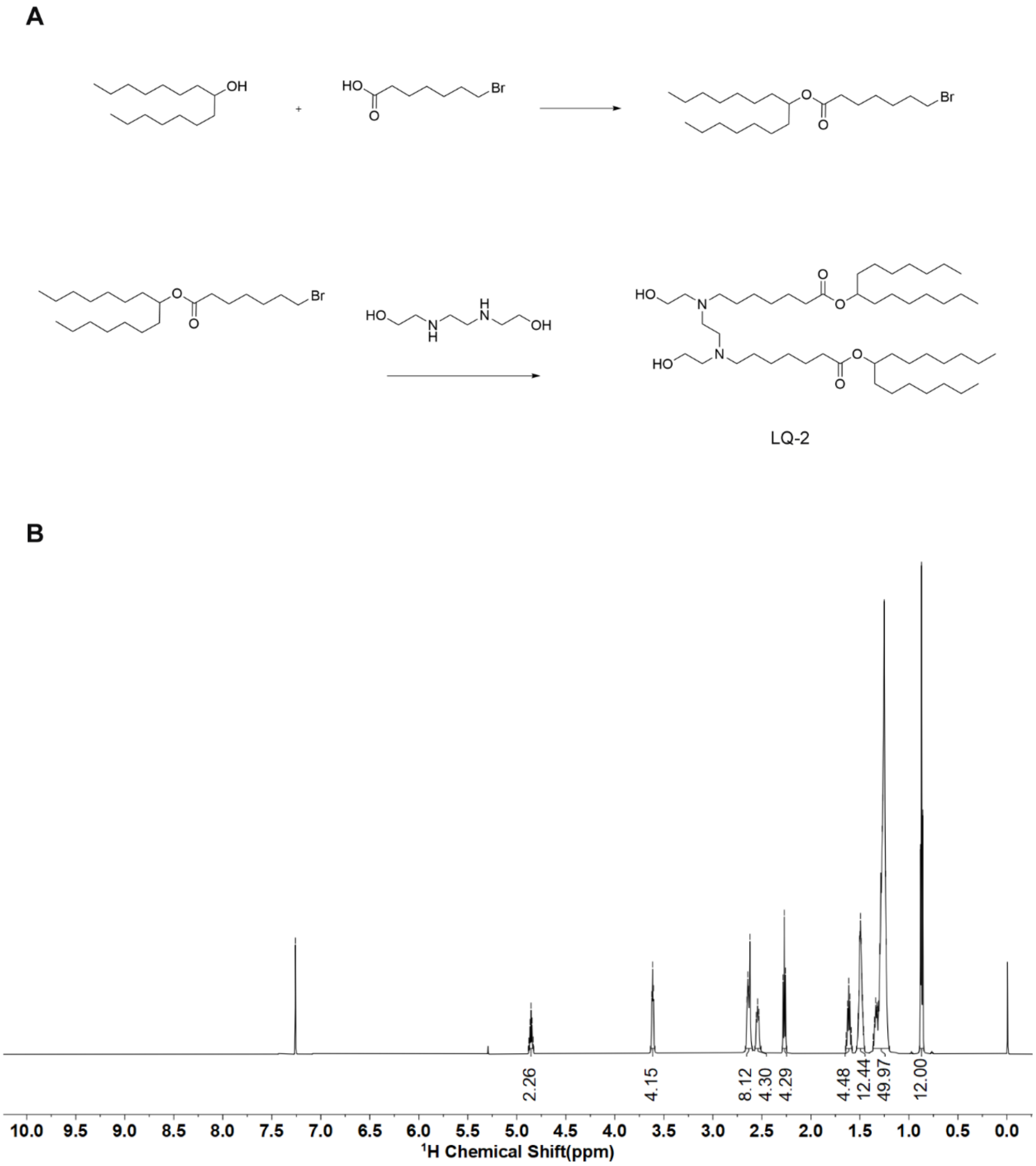


**Figure S2. The synthetic route (A) and $^1$H NMR (B) of LQ-2.** $^1$H NMR (600 MHz, $CDCl_3$) δ ： 4.85 (p, *J* = 6.2 Hz, 2H), 3.61 (t, *J* = 4.9 Hz, 4H), 2.67 – 2.60 (m, 8H), 2.54 (t, *J* = 7.9 Hz, 4H), 2.27 (t, *J* = 7.5 Hz, 4H), 1.61 (p, *J* = 7.5 Hz, 4H), 1.50 (qd, *J* = 7.6, 3.4 Hz, 12H), 1.37 – 1.20 (m, 50H), 0.87 (t, *J* = 7.0 Hz, 12H).

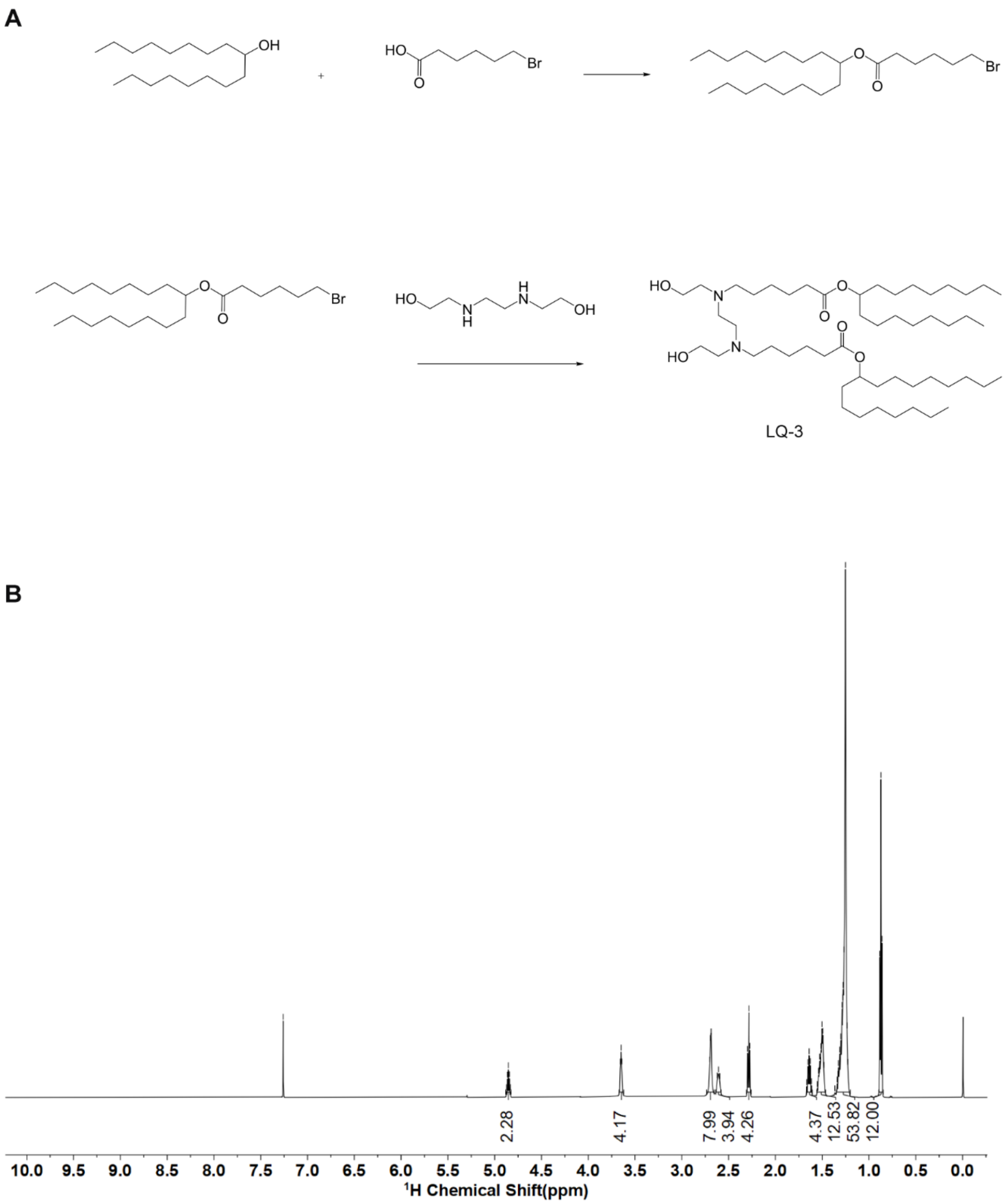


**Figure S3. The synthetic route (A) and $^1$H NMR (B) of LQ-3.** $^1$H NMR (600 MHz, $CDCl_3$) δ :4.85 (p, *J* = 6.2 Hz, 2H), 3.65 (t, *J* = 4.8 Hz, 4H), 2.73 – 2.65 (m, 8H), 2.61 (t, *J* = 8.1 Hz, 4H), 2.28 (t, *J* = 7.4 Hz, 4H), 1.64 (p, *J* = 7.5 Hz, 4H), 1.51 (dq, *J* = 19.0, 7.3, 6.8 Hz, 12H), 1.35 – 1.20 (m, 54H), 0.87 (t, *J* = 6.9 Hz, 12H).

A

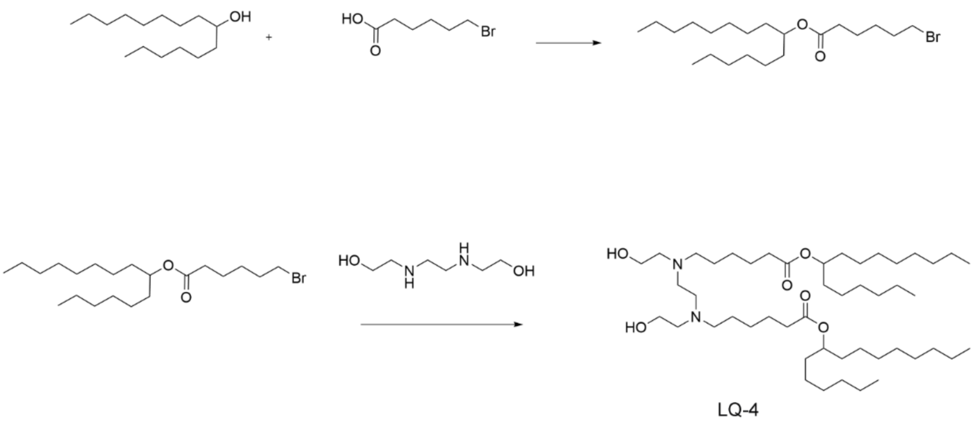


B

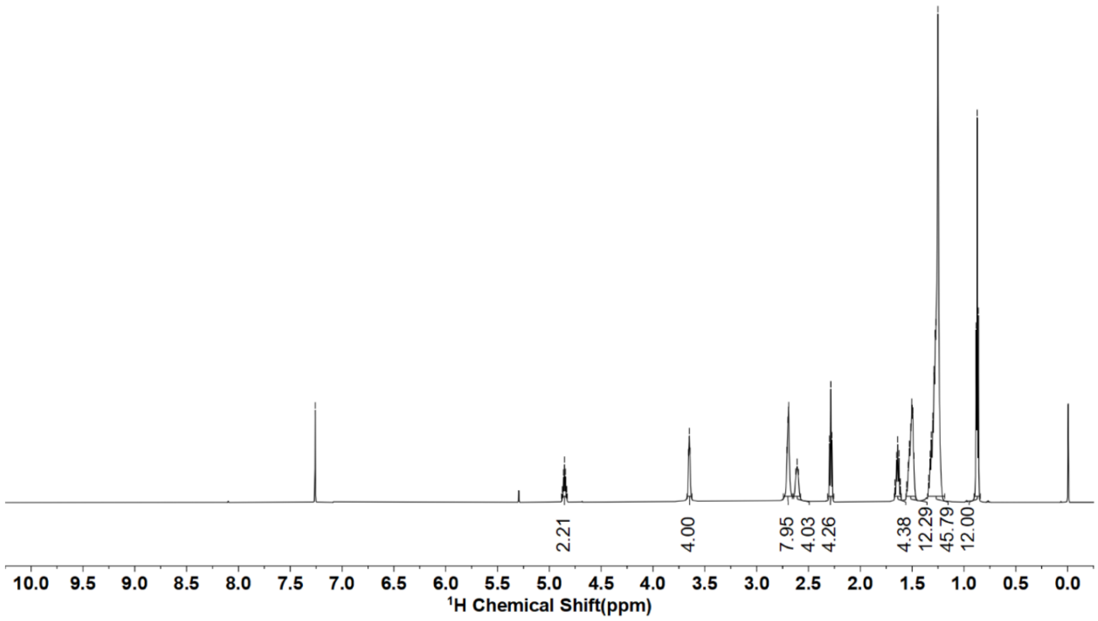


**Figure S4. The synthetic route (A) and $^1$H NMR (B) of LQ-4.** $^1$H NMR (600 MHz, $CDCl_3$) δ： 4.85 (p, *J* = 6.3 Hz, 2H), 3.65 (t, *J* = 4.8 Hz, 4H), 2.69 (d, *J* = 5.2 Hz, 8H), 2.62 (d, *J* = 8.2 Hz, 4H), 2.28 (t, *J* = 7.4 Hz, 4H), 1.64 (p, *J* = 7.6 Hz, 4H), 1.51 (dq, *J* = 18.9, 6.9, 6.0 Hz, 12H), 1.35 – 1.18 (m, 46H), 0.87 (t, *J* = 6.9 Hz, 12H).

A

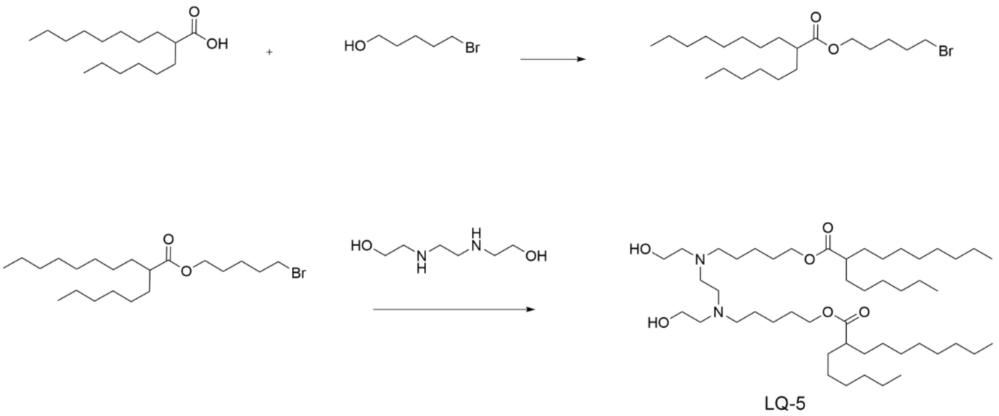

B

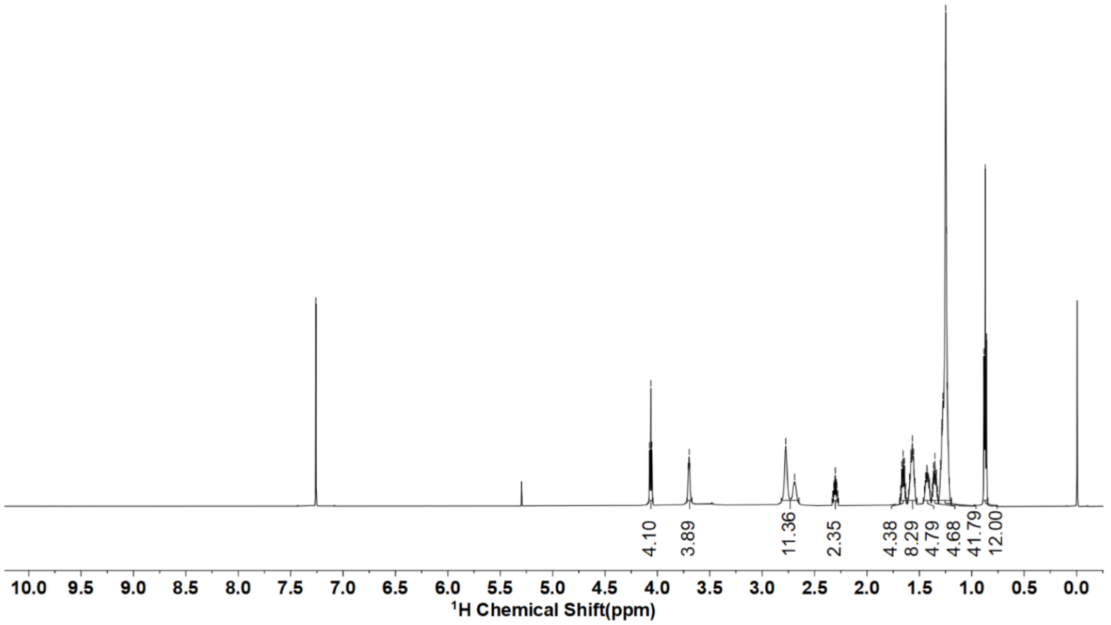

**Figure S5. The synthetic route (A) and $^1$H NMR (B) of LQ-5.** $^1$H NMR (600 MHz, $CDCl_3$) δ：4.06 (t, *J* = 6.6 Hz, 4H), 3.70 (t, *J* = 4.8 Hz, 4H), 2.73 (d, *J* = 51.1 Hz, 12H), 2.30 (tt, *J* = 8.8, 5.3 Hz, 2H), 1.66 (p, *J* = 6.9 Hz, 4H), 1.57 (p, *J* = 7.9 Hz, 8H), 1.46 – 1.39 (m, 4H), 1.35 (p, *J* = 7.5 Hz, 4H), 1.32 – 1.19 (m, 42H), 0.87 (t, *J* = 6.9 Hz, 12H).

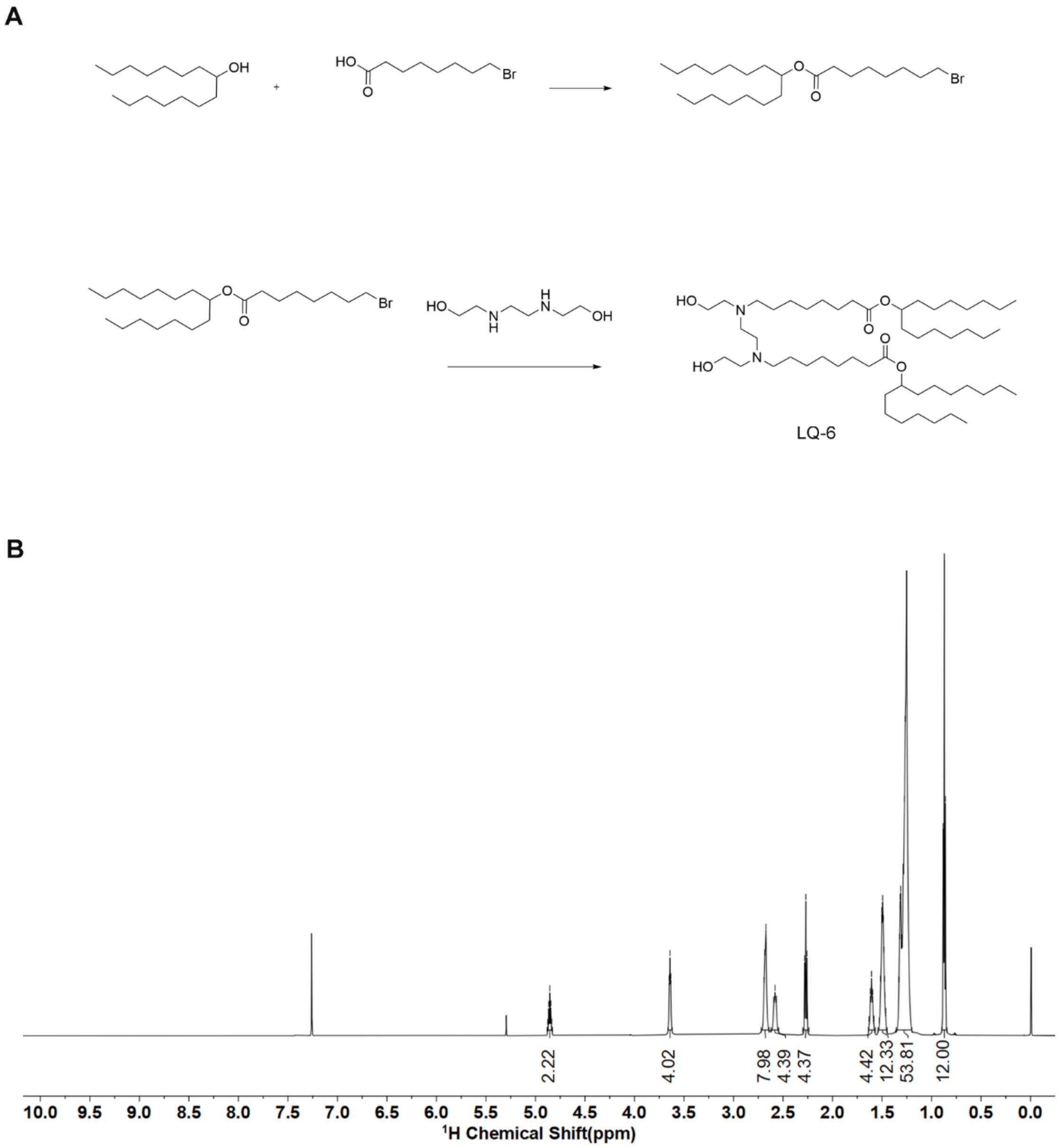


**Figure S6. The synthetic route (A) and $^1$H NMR (B) of LQ-6.** $^1$H NMR (600 MHz, $CDCl_3$) δ: 4.86 (p, *J* = 6.2 Hz, 2H), 3.64 (t, *J* = 4.8 Hz, 4H), 2.68 (d, *J* = 6.8 Hz, 8H), 2.58 (t, *J* = 8.0 Hz, 4H), 2.27 (t, *J* = 7.5 Hz, 4H), 1.61 (p, *J* = 7.2 Hz, 4H), 1.49 (qd, *J* = 7.7, 5.2, 4.2 Hz, 12H), 1.36 – 1.20 (m, 54H), 0.87 (t, *J* = 7.0 Hz, 12H).

**A**

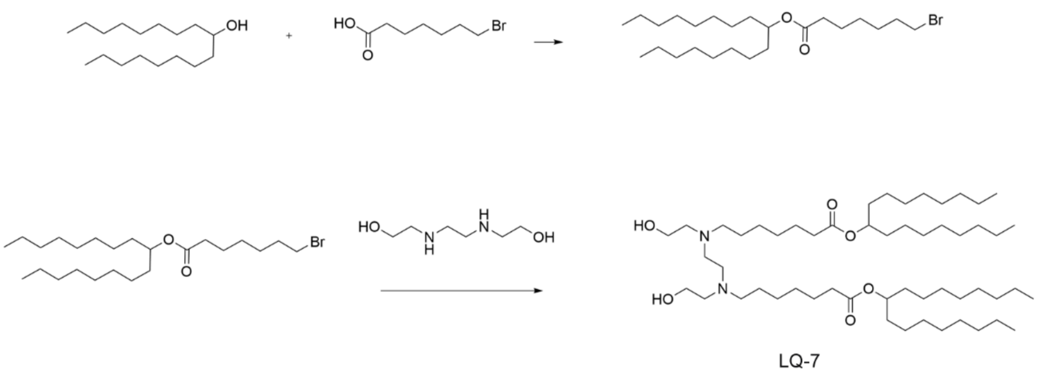


**B**

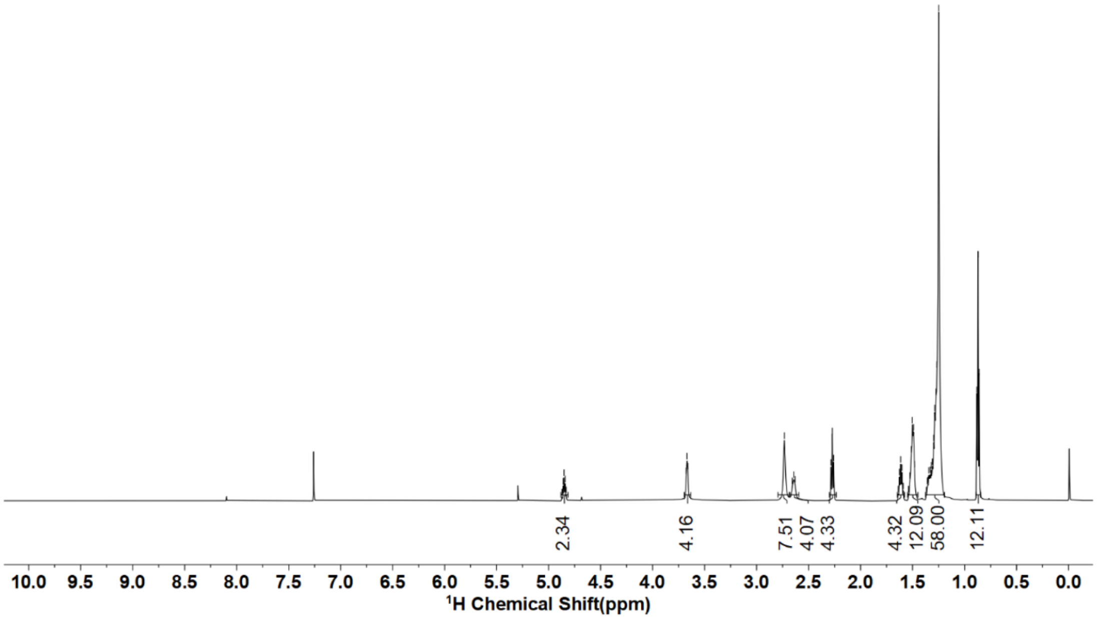


**Figure S7. The synthesis route (A) and $^1$H NMR (B) of LQ-7.** $^1$H NMR (600 MHz, $CDCl_3$) δ： 4.85 (p, *J* = 6.2 Hz, 2H), 3.67 (t, *J* = 4.9 Hz, 4H), 2.73 (s, 8H), 2.64 (t, *J* = 8.0 Hz, 4H), 2.27 (t, *J* = 7.5 Hz, 4H), 1.61 (p, *J* = 7.4 Hz, 4H), 1.51 (dd, *J* = 13.6, 7.0 Hz, 12H), 1.38 – 1.19 (m, 58H), 0.87 (t, *J* = 7.0 Hz, 12H).

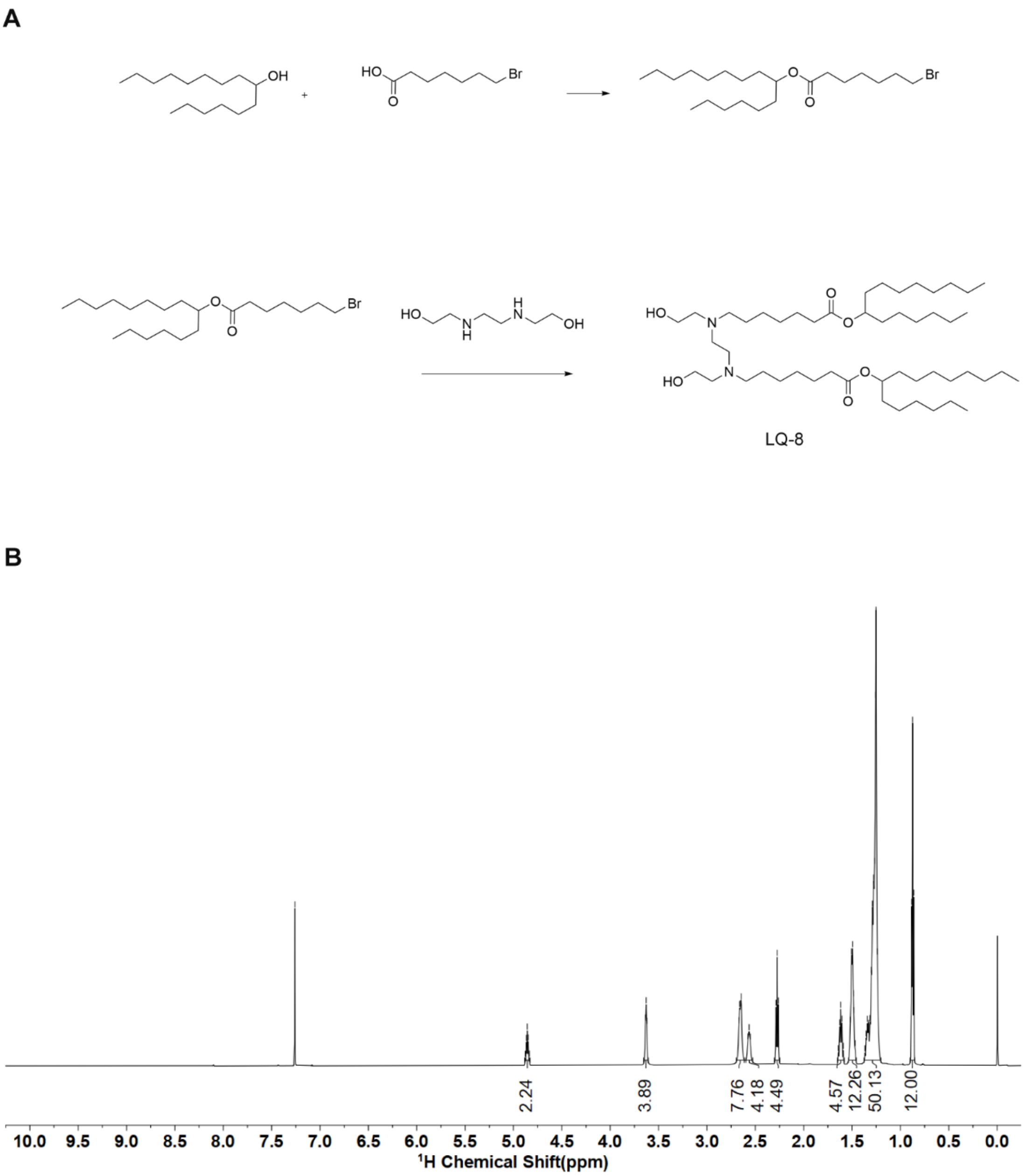


**Figure S8. The synthesis route (A) and $^1$H NMR (B) of LQ-8.** $^1$H NMR (600 MHz, $CDCl_3$) δ：4.86 (p, *J* = 6.3 Hz, 2H), 3.63 (t, *J* = 4.8 Hz, 4H), 2.66 (dd, *J* = 9.9, 5.1 Hz, 8H), 2.56 (t, *J* = 8.0 Hz, 4H), 2.27 (t, *J* = 7.4 Hz, 4H), 1.62 (p, *J* = 7.5 Hz, 4H), 1.49 (dd, *J* = 10.3, 4.9 Hz, 12H), 1.38 – 1.20 (m, 50H), 0.87 (t, *J* = 7.0 Hz, 12H).

A

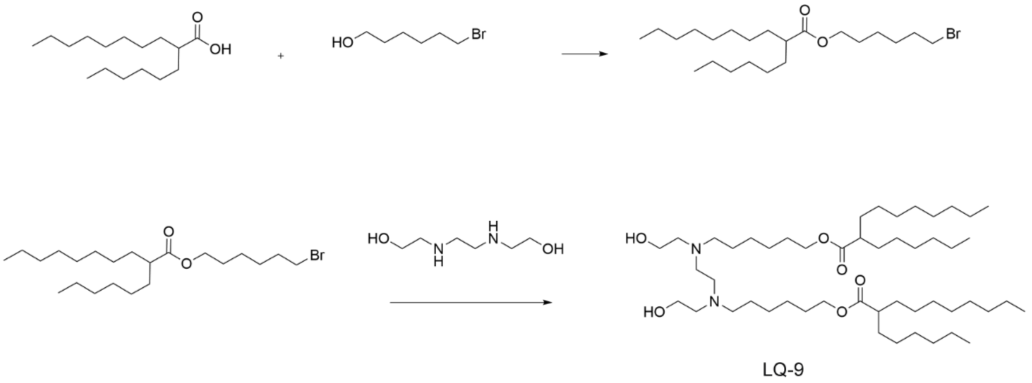


B

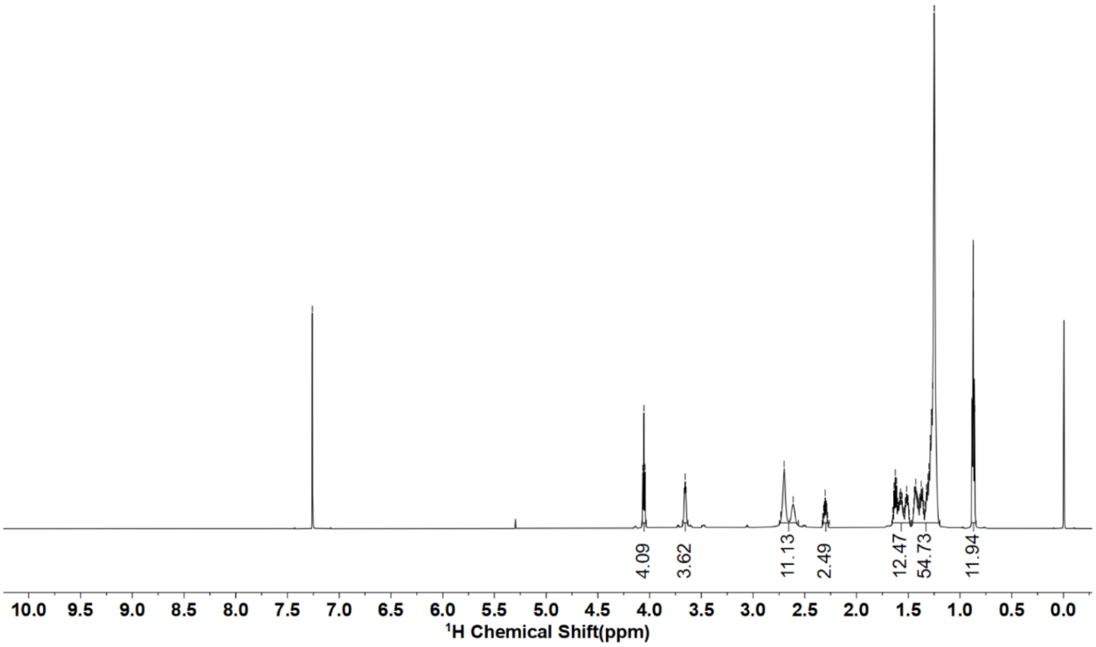


**Figure S9. The synthetic route (A) and $^1$H NMR (B) of LQ-9.** $^1$H NMR (600 MHz, $CDCl_3$) δ： 4.05 (t, *J* = 6.6 Hz, 4H), 3.66 (t, *J* = 4.8 Hz, 4H), 2.66 (d, *J* = 52.0 Hz, 12H), 2.30 (tt, *J* = 8.9, 5.3 Hz, 2H), 1.66 – 1.48 (m, 12H), 1.46 – 1.20 (m, 54H), 0.87 (td, *J* = 7.0, 1.5 Hz, 12H).

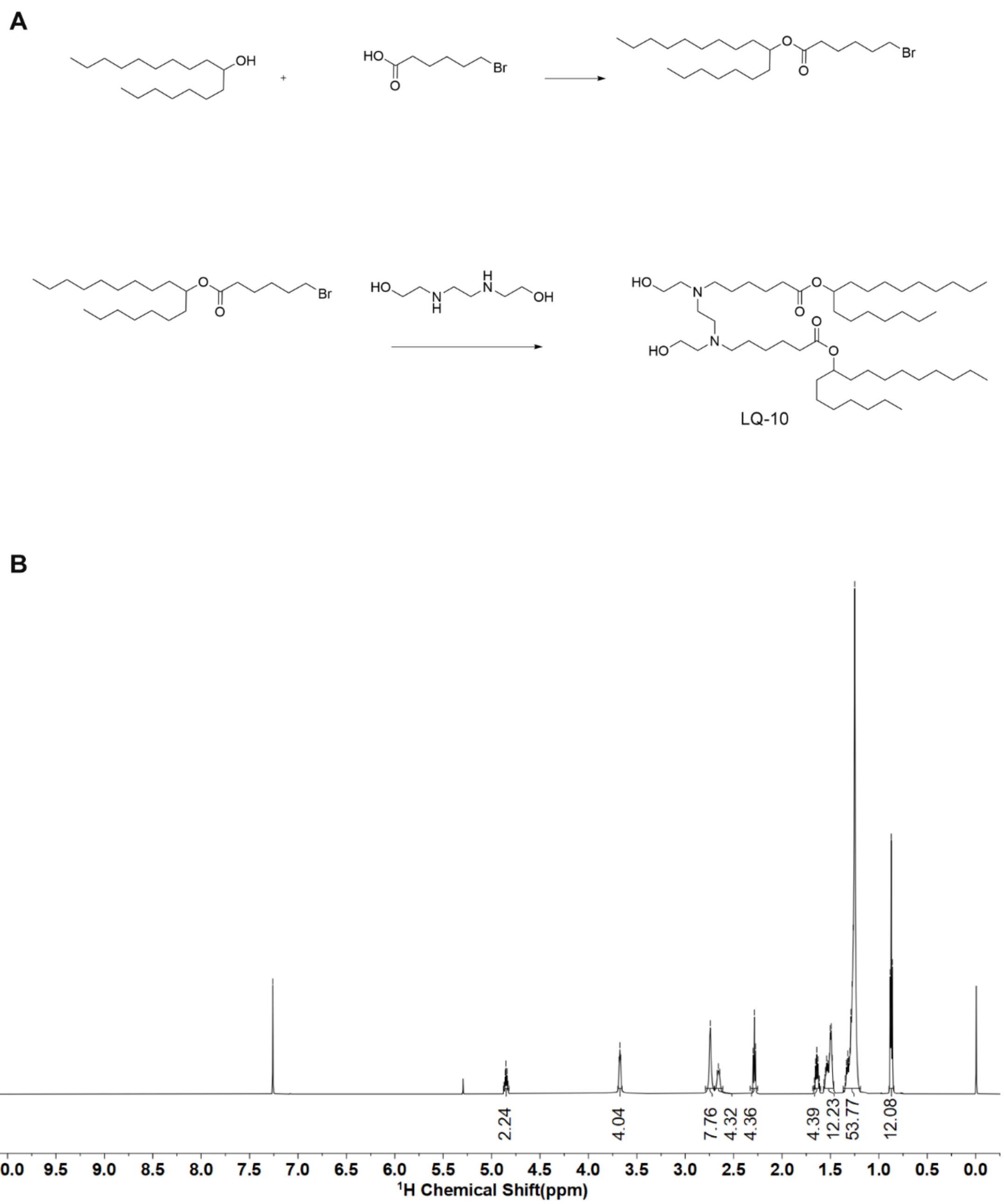


**Figure S10. The synthetic route (A) and $^1$H NMR (B) of LQ-10.** $^1$H NMR (600 MHz, $CDCl_3$) δ: 4.85 (p, $J$ = 6.3 Hz, 2H), 3.67 (t, $J$ = 4.8 Hz, 4H), 2.74 (s, 8H), 2.66 (t, $J$ = 8.0 Hz, 4H), 2.29 (t, $J$ = 7.4 Hz, 4H), 1.64 (p, $J$ = 7.5 Hz, 4H), 1.52 (dq, $J$ = 26.5, 6.9, 6.0 Hz, 12H), 1.37 – 1.19 (m, 54H), 0.87 (t, $J$ = 6.9 Hz, 12H).

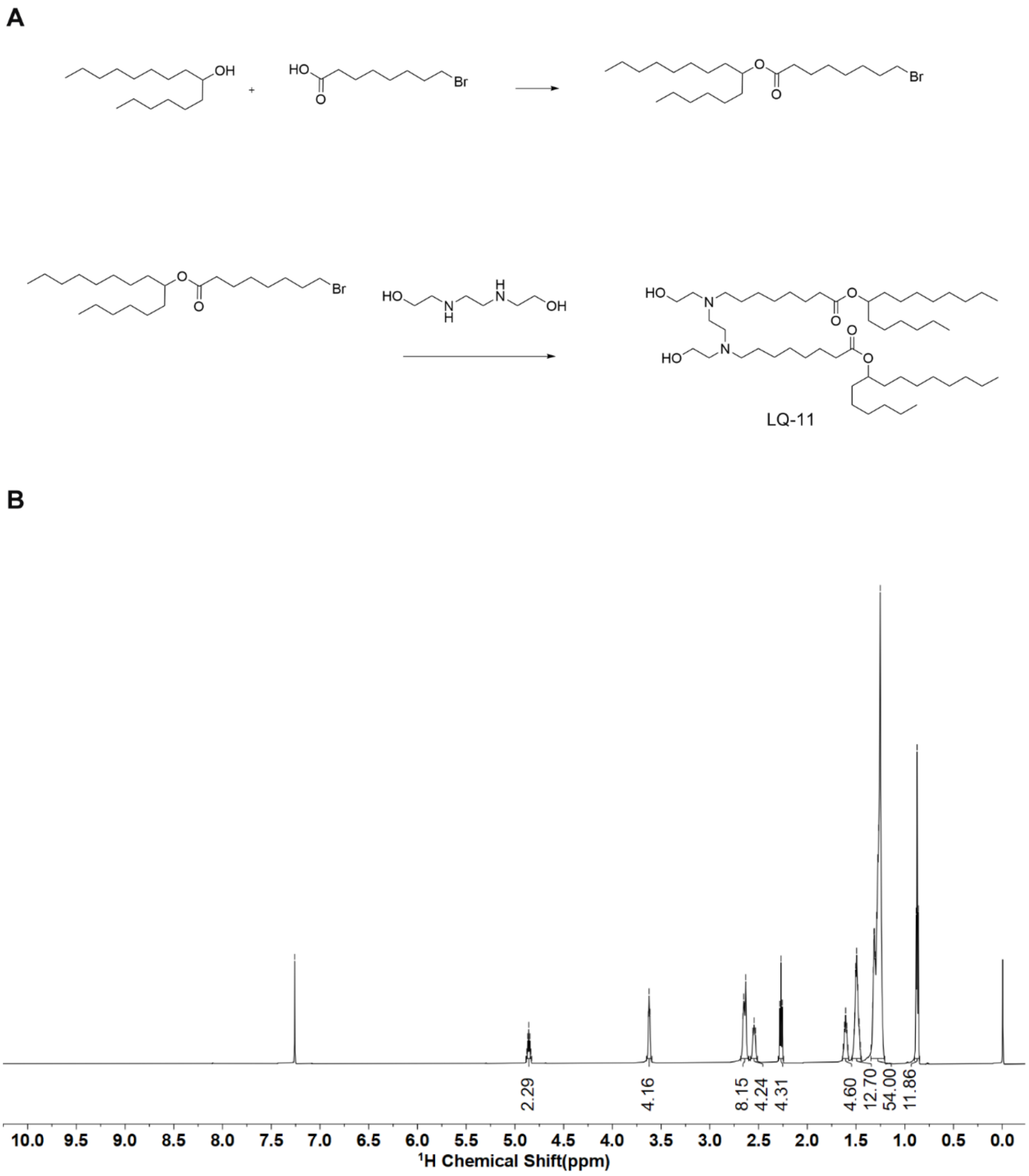


**Figure S11. The synthetic route (A) and $^1$H NMR (B) of LQ-11.** $^1$H NMR (600 MHz, $CDCl_3$) δ: 4.86 (p, *J* = 6.3 Hz, 2H), 3.62 (t, *J* = 4.9 Hz, 4H), 2.68 – 2.60 (m, 8H), 2.55 (t, *J* = 8.0 Hz, 4H), 2.27 (t, *J* = 7.5 Hz, 4H), 1.61 (t, *J* = 7.4 Hz, 4H), 1.49 (p, *J* = 7.7, 6.7 Hz, 12H), 1.35 – 1.21 (m, 54H), 0.87 (t, *J* = 7.0 Hz, 12H).

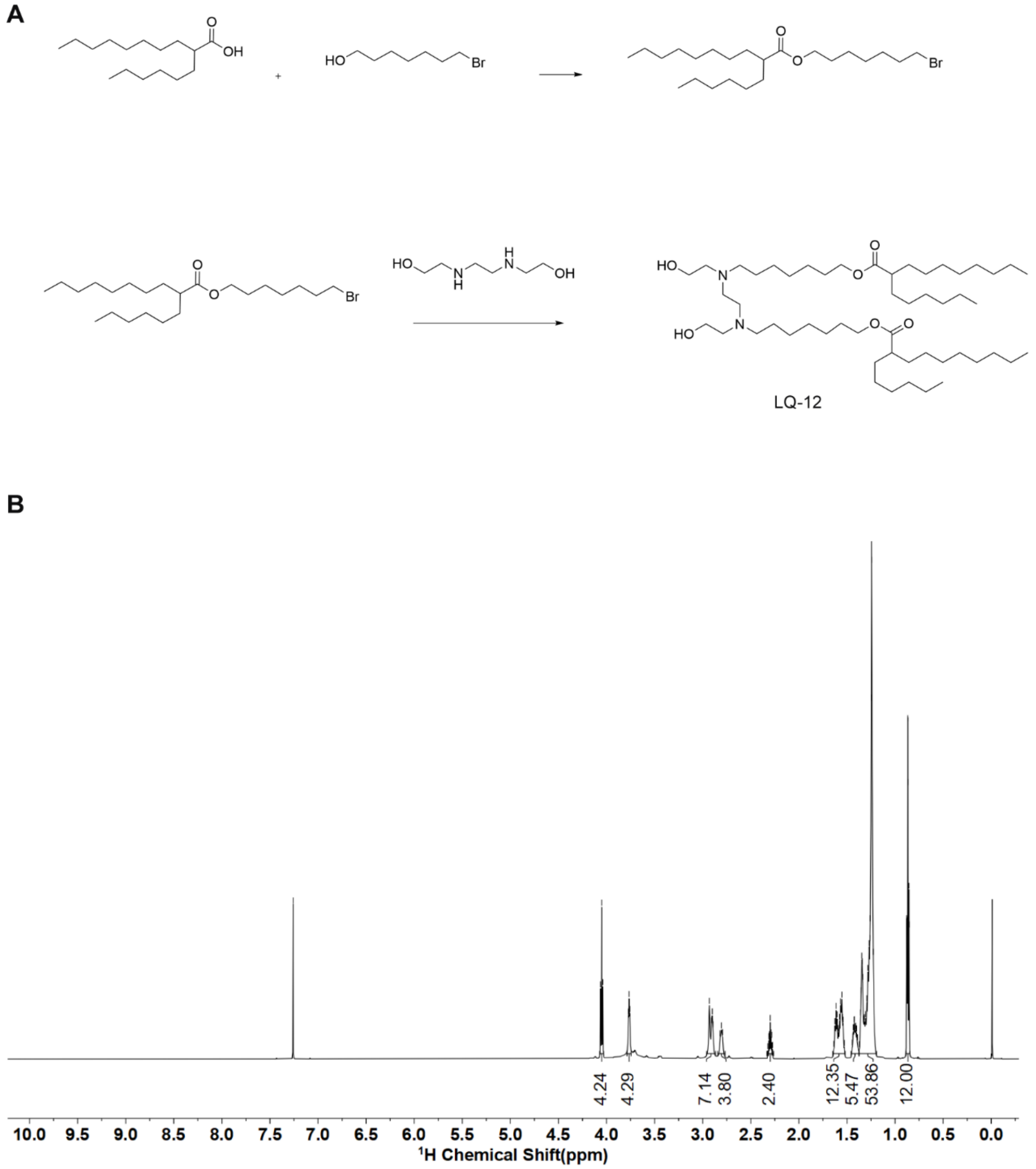


**Figure S12. The synthetic route (A) and $^1$H NMR (B) of LQ-12.** $^1$H NMR (600 MHz, $CDCl_3$) δ: 4.05 (t, $J$ = 6.7 Hz, 4H), 3.77 (t, $J$ = 4.8 Hz, 4H), 2.96 – 2.87 (m, 8H), 2.81 (t, $J$ = 8.2 Hz, 4H), 2.30 (tt, $J$ = 8.9, 5.3 Hz, 2H), 1.65 – 1.52 (m, 12H), 1.46 – 1.38 (m, 4H), 1.37 – 1.19 (m, 54H), 0.87 (td, $J$ = 7.1, 1.5 Hz, 12H).

A

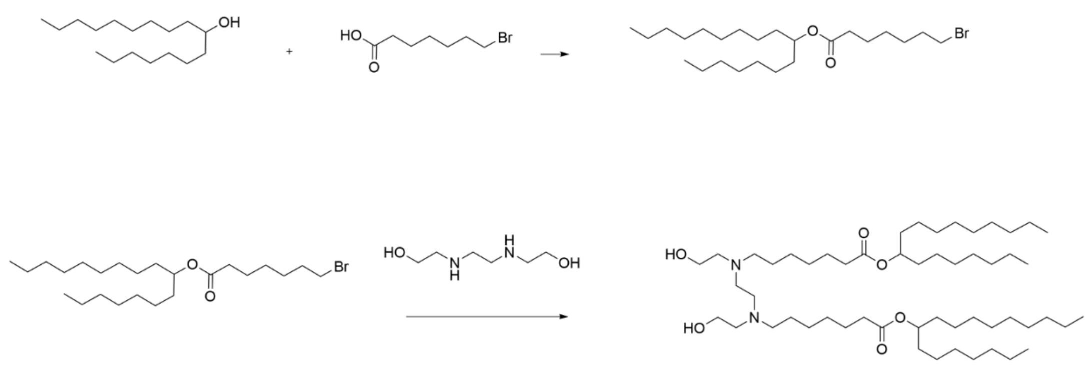


B

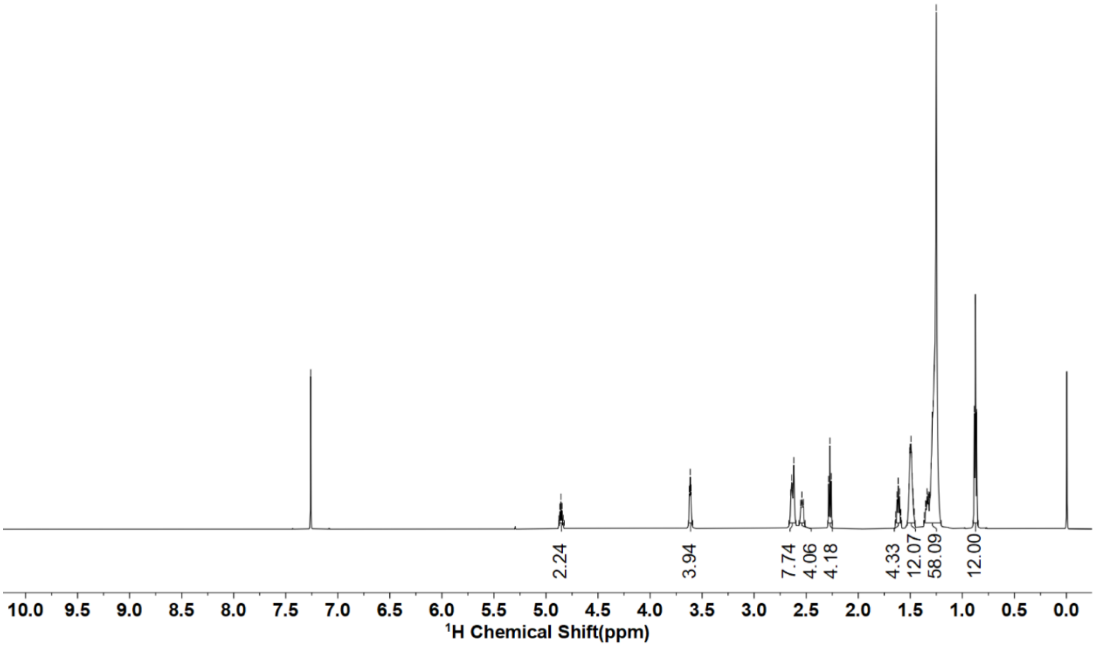


**Figure S13. The synthetic route (A) and $^1$H NMR (B) of LQ-13.** $^1$H NMR (600 MHz, $CDCl_3$) δ: 4.86 (p, $J$ = 6.2 Hz, 2H), 3.61 (t, $J$ = 4.8 Hz, 4H), 2.67 – 2.60 (m, 8H), 2.54 (t, $J$ = 8.0 Hz, 4H), 2.27 (t, $J$ = 7.5 Hz, 4H), 1.62 (p, $J$ = 7.5 Hz, 4H), 1.49 (tt, $J$ = 7.6, 4.6 Hz, 12H), 1.37 – 1.20 (m, 58H), 0.87 (t, $J$ = 6.9 Hz, 12H).

**A**

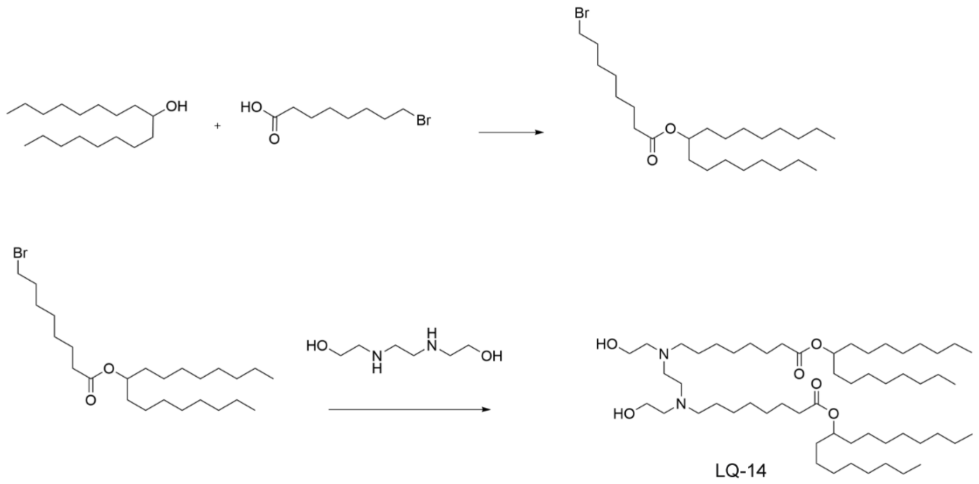


**B**

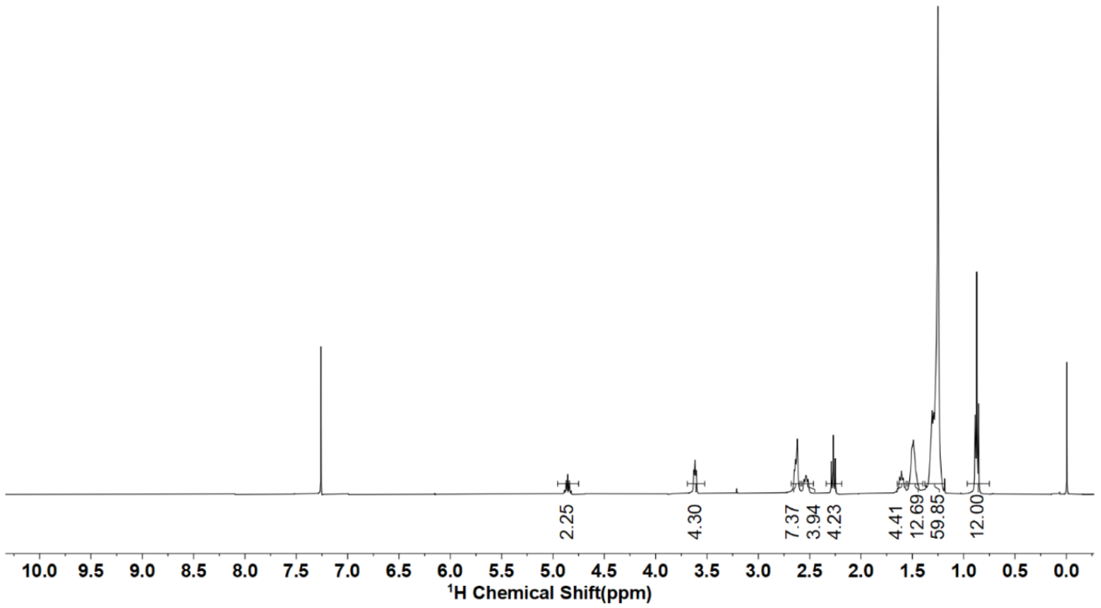


**Figure S14. The synthetic route (A) and $^1$H NMR (B) of LQ-14.** $^1$H NMR (400 MHz, $CDCl_3$) δ： 4.96 – 4.75 (m, 2H), 3.69 – 3.52 (m, 4H), 2.68 – 2.59 (m, 8H), 2.58 – 2.46 (m, 4H), 2.27 (t, *J* = 7.5 Hz, 4H), 1.65 – 1.56 (m, 4H), 1.49 (s, 12H), 1.28 (d, *J* = 22.1 Hz, 60H), 0.87 (t, *J* = 6.8 Hz, 12H).

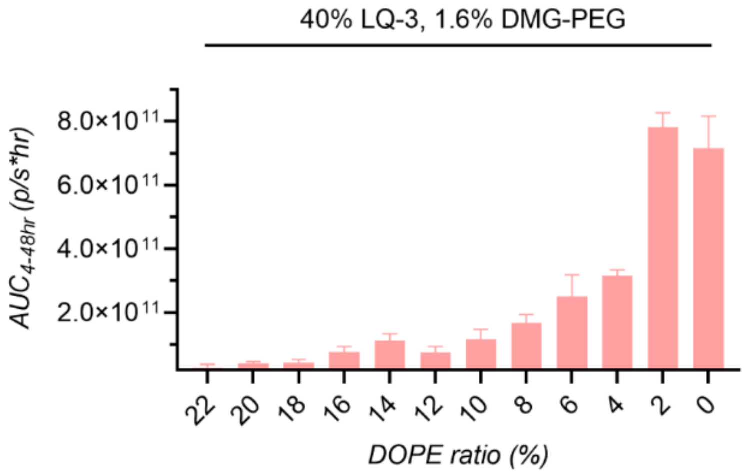


**Figure S15. Determination of the optimal DOPE ratio in formulations.** Area under the curve (AUC) for whole-body luciferase bioluminescence signals from LQ-3 LNPs with varying DOPE ratios post intravenous administration at an mRNA dosage of 0.25 mg/kg (n = 3, mean ± SD).

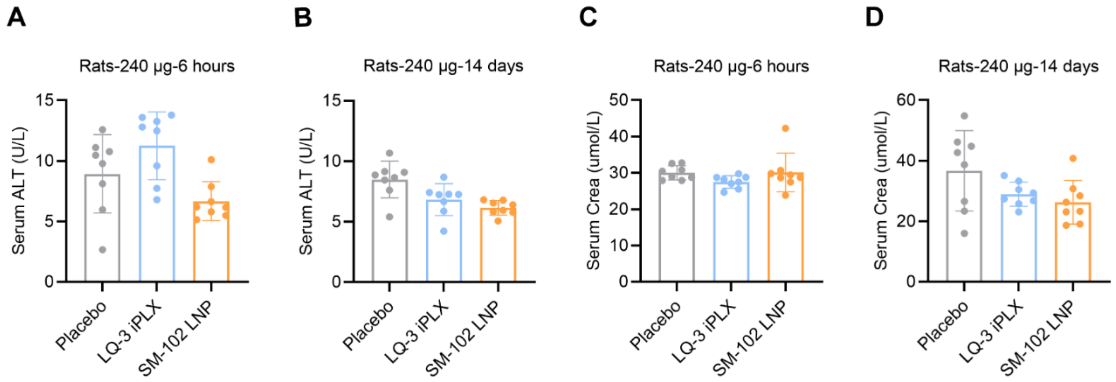


**Figure S16. In vivo safety assessment in rats: Serum ALT and creatinine levels.** Serum concentrations of ALT (A, B) and creatinine (C, D) in SD rats recorded at 6 hours and 14 days post intramuscular administration. Sample size: n = 8 (comprising 4 females and 4 males); presented as mean ± SD. No notable difference was found between the placebo and mRNA-LNP groups when administered at 240 µg/injection.

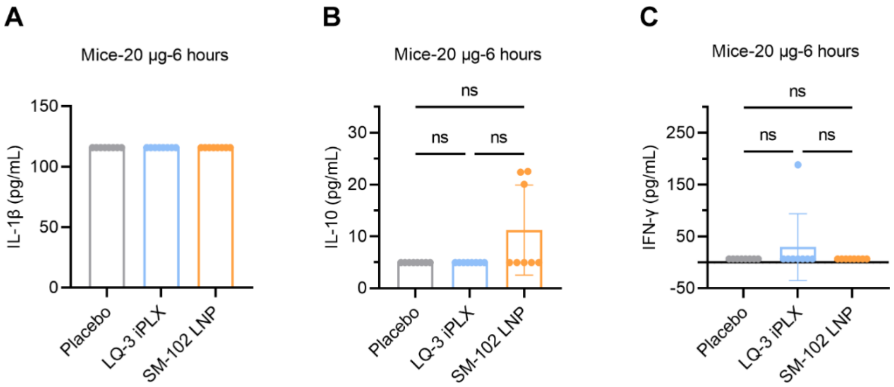


**Figure S17. In vivo safety assessment in mice: Serum cytokine levels.** Serum concentrations of cytokines IL-1β (A), IL-10 (B), and IFN-γ (C) were measured in mice 6 hours post intramuscular administration of mRNA-LNP at a dosage of 20 μg/injection. Sample size: n = 8 (comprising 4 females and 4 males); data is represented as mean ± SD. No notable difference was detected between the placebo and mRNA-LNP groups when administered at 20 μg/injection.

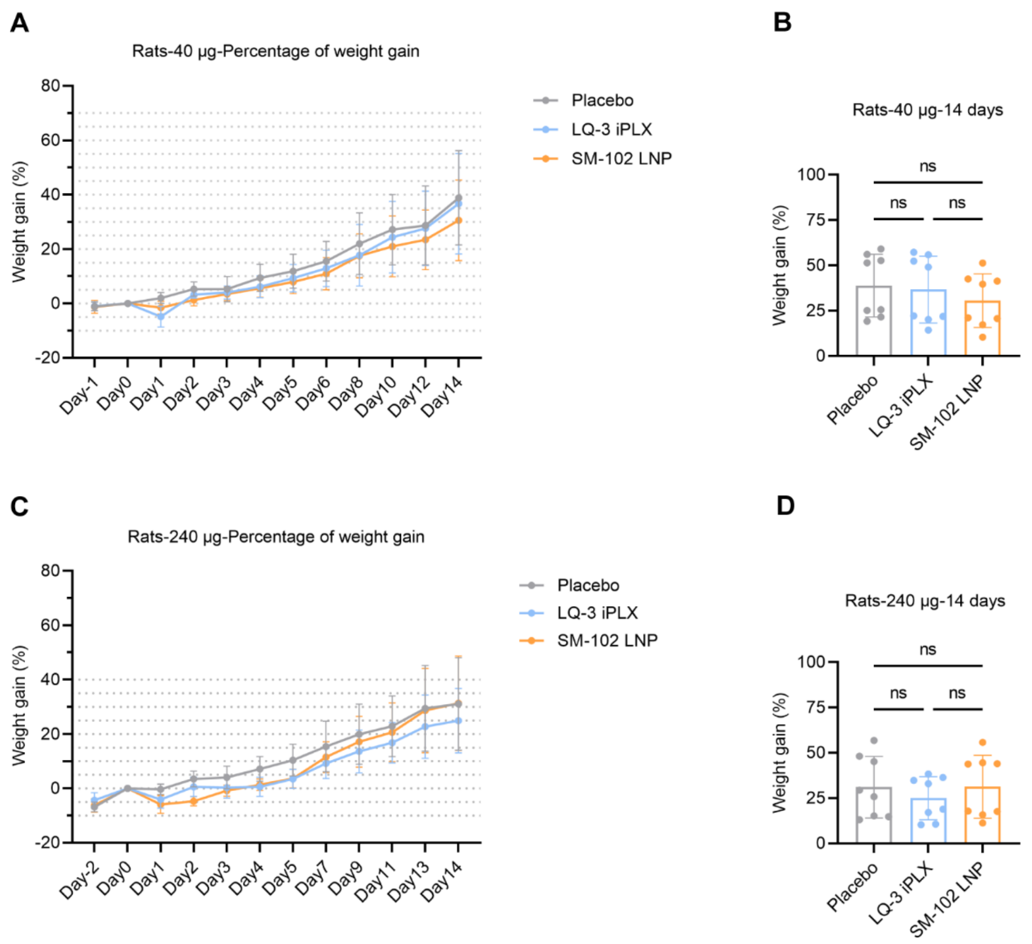


**Figure S18. Body weight progression during *in vivo* safety assessment in rats.** (A) Rat body weight percentage increase post intramuscular injection of mRNA-LNP at 40 µg/injection over time, starting from Day 0 as the baseline. (B) Percentage body weight gain on Day 14 after the 40 µg/injection of mRNA-LNP. (C) Rat body weight percentage increase following mRNA-LNP administration at 240 µg/injection, with Day 0 as the reference. (D) Weight gain percentage on Day 14 after the 240 µg/injection. Across all data points, no marked differences were noted between the placebo and mRNA-LNP groups. The study comprised 8 rats, split evenly between females and males. Data is presented as mean ± SD.

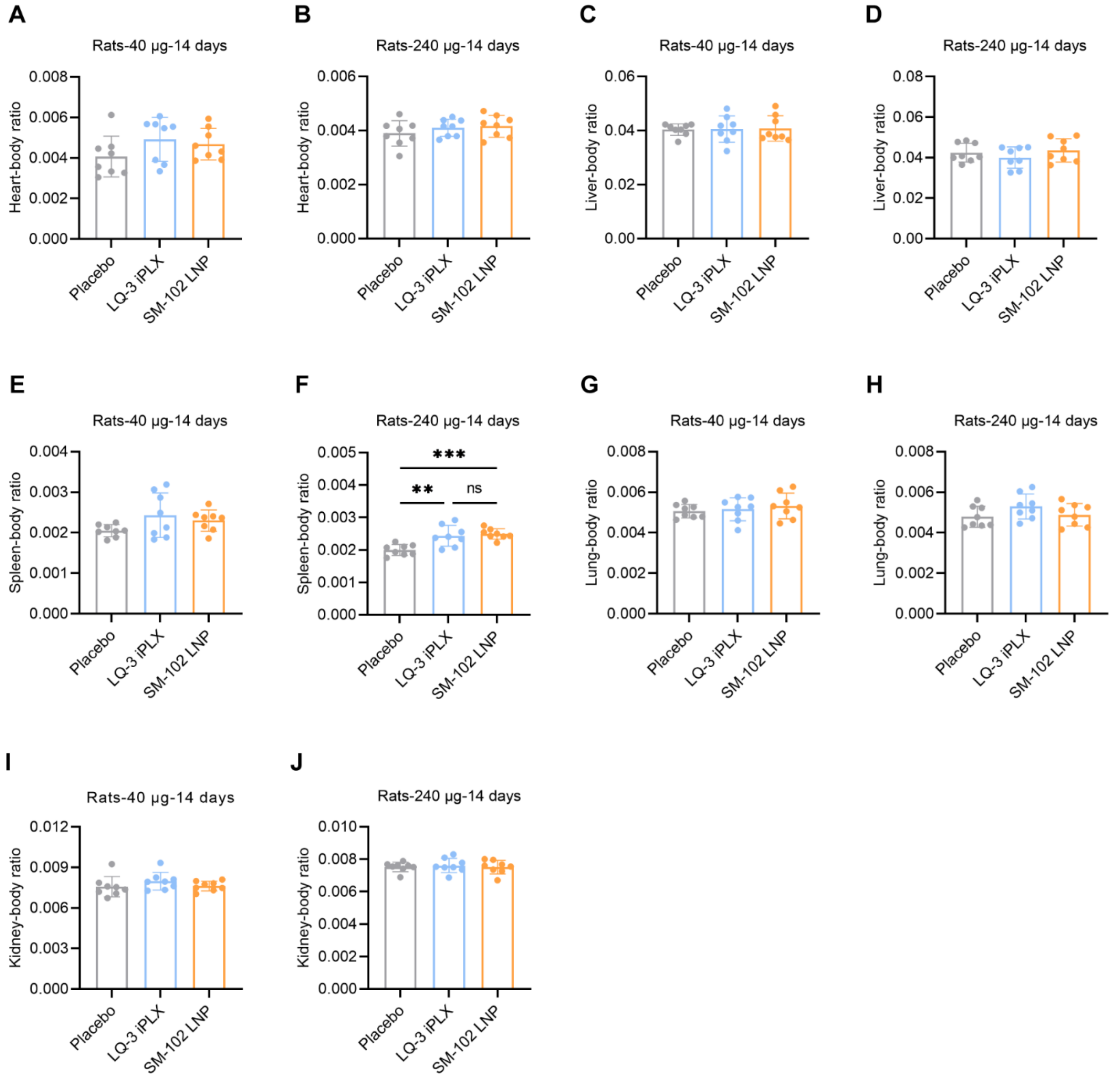


**Figure S19. Ratios of organ-to-body weight post mRNA-LNP administration.** After 14 days of intramuscular injection of mRNA-LNP at doses of 40 µg/injection and 240 µg/injection, organ-to-body weight ratios were determined for the heart (A, B), liver (C, D), spleen (E, F), lungs (G, H), and kidneys (I, J). The spleen-to-body weight ratio showed a significant difference at the higher dose of 240 µg/injection; however, the other organs showed no noticeable difference between the placebo and mRNA-LNP groups. The study included 8 rats, equally divided between females and males. Data is depicted as mean ± SD. $^{**}p \leq 0.01$, $^{***}p \leq 0.001$.

**Table S1.** Physicochemical properties of formulations with varied DSPC ratios.

| LNP | Group | % mole ratios (ionizable lipids/DSPC/Cholesterol/DMG-PEG2k) | N/P | Size (nm) | PDI | EE (%) |
|---|---|---|---|---|---|---|
| 1 | 0% DSPC | 40：0：58.4：1.6 | 8 | 90.1 | 0.139 | 97.1 |
| 2 | 2% DSPC | 40：2：56.4：1.6 | 8 | 82.5 | 0.081 | 96.8 |
| 3 | 4% DSPC | 40：4：54.4：1.6 | 8 | 73.3 | 0.073 | 97.3 |
| 4 | 6% DSPC | 40：6：52.4：1.6 | 8 | 70.8 | 0.101 | 97.3 |
| 5 | 8% DSPC | 40：8：50.4：1.6 | 8 | 66.6 | 0.100 | 97.1 |
| 6 | 10% DSPC | 40：10：48.4：1.6 | 8 | 69.0 | 0.092 | 96.9 |
| 7 | 12% DSPC | 40：12：46.4：1.6 | 8 | 66.0 | 0.063 | 97.2 |
| 8 | 14% DSPC | 40：14：44.4：1.6 | 8 | 63.8 | 0.072 | 97.0 |
| 9 | 16% DSPC | 40：16：42.4：1.6 | 8 | 69.9 | 0.089 | 97.0 |
| 10 | 18% DSPC | 40：18：40.4：1.6 | 8 | 64.3 | 0.103 | 97.2 |
| 11 | 20% DSPC | 40：20：38.4：1.6 | 8 | 66.0 | 0.073 | 97.2 |
| 12 | 22% DSPC | 40：22：36.4：1.6 | 8 | 63.3 | 0.094 | 96.8 |
| 13 | 24% DSPC | 40：24：34.4：1.6 | 8 | 63.6 | 0.053 | 97.8 |
| 14 | 26% DSPC | 40：26：32.4：1.6 | 8 | 70.5 | 0.105 | 97.4 |
| 15 | 28% DSPC | 40：28：30.4：1.6 | 8 | 71.0 | 0.099 | 97.5 |
| 16 | 30% DSPC | 40：30：28.4：1.6 | 8 | 90.8 | 0.105 | 97.5 |
| 17 | 32% DSPC | 40：32：26.4：1.6 | 8 | 79.6 | 0.113 | 97.2 |
| 18 | 34% DSPC | 40：34：24.4：1.6 | 8 | 92.2 | 0.121 | 97.0 |
| 19 | 36% DSPC | 40：36：22.4：1.6 | 8 | 84.9 | 0.145 | 97.6 |
| 20 | 38% DSPC | 40：38：20.4：1.6 | 8 | 100.2 | 0.157 | 97.0 |
| 21 | 40% DSPC | 40：40：18.4：1.6 | 8 | 90.6 | 0.166 | 97.5 |

**Table S2.** Physicochemical properties of formulations based on varying ionizable lipid and DSPC ratios.

| LNP | Group | % mole ratios (ionizable lipids/DSPC/Cholesterol/ DMG-PEG2k) | N/P | Size (nm) | PDI | EE (%) |
|---|---|---|---|---|---|---|
| 1 | 35% LQ-3 & 0% DSPC | 35：0：63.4：1.6 | 8 | 81.6 | 0.206 | 94.9 |
| 2 | 40% LQ-3& 0% DSPC | 40：0：58.4：1.6 | 8 | 87.1 | 0.159 | 96.0 |
| 3 | 45% LQ-3 & 0% DSPC | 45：0：53.4：1.6 | 8 | 101.3 | 0.137 | 95.9 |
| 4 | 30% LQ-3 & 2% DSPC | 30：2：66.4：1.6 | 8 | 65.4 | 0.197 | 94.8 |
| 5 | 35% LQ-3 & 2% DSPC | 35：2：61.4：1.6 | 8 | 73.4 | 0.180 | 95.8 |
| 6 | 40% LQ-3 & 2% DSPC | 40：2：56.4：1.6 | 8 | 85.8 | 0.159 | 96.3 |
| 7 | 45% LQ-3 & 2% DSPC | 45：2：51.4：1.6 | 8 | 92.4 | 0.142 | 96.4 |
| 8 | 50% LQ-3 & 2% DSPC | 50：2：46.4：1.6 | 8 | 101.7 | 0.109 | 96.4 |
| 9 | 60% LQ-3 & 2% DSPC | 60：2：36.4：1.6 | 8 | 121.6 | 0.050 | 95.6 |
| 10 | 25% LQ-3 & 4% DSPC | 25：4：69.4：1.6 | 8 | 72.9 | 0.287 | 95.3 |
| 11 | 30% LQ-3 & 4% DSPC | 30：4：64.4：1.6 | 8 | 61.0 | 0.174 | 96.0 |
| 12 | 35% LQ-3 & 4% DSPC | 35：4：59.4：1.6 | 8 | 65.3 | 0.156 | 96.1 |
| 13 | 40% LQ-3 & 4% DSPC | 40：4：54.4：1.6 | 8 | 70.3 | 0.180 | 96.1 |
| 14 | 45% LQ-3 & 4% DSPC | 45：4：49.4：1.6 | 8 | 76.6 | 0.154 | 97.3 |
| 15 | 60% LQ-3 & 4% DSPC | 60：4：34.4：1.6 | 8 | 115.0 | 0.056 | 95.1 |
| 16 | 25% LQ-3 & 10% DSPC | 25：10：63.4：1.6 | 8 | 59.0 | 0.209 | 97.6 |
| 17 | 30% LQ-3 & 10% DSPC | 30：10：58.4：1.6 | 8 | 55.4 | 0.155 | 97.2 |
| 18 | 35% LQ-3 & 10% DSPC | 35：10：53.4：1.6 | 8 | 56.5 | 0.182 | 97.0 |
| 19 | 40% LQ-3 & 10% DSPC | 40：10：48.4：1.6 | 8 | 63.9 | 0.180 | 97.6 |
| 20 | 45% LQ-3 & 10% DSPC | 45：10：43.4：1.6 | 8 | 68.3 | 0.151 | 97.0 |

**Table S3.** Physicochemical Properties of Formulations Varying in PEG-Lipid Ratios.

| LNP | Group | % mole ratios (ionizable lipids/DSPC/Cholesterol/DMG-PEG2k) | N/P | Size (nm) | PDI | EE (%) |
|---|---|---|---|---|---|---|
| 1 | 45% LQ-3 & 0% DSPC &1.6% PEG | 45：0：53.4：1.6 | 8 | 109.1 | 0.119 | 98.2 |
| 2 | 45% LQ-3 & 0% DSPC & 2% PEG | 45：0：53：2 | 8 | 103.8 | 0.113 | 97.6 |
| 3 | 45% LQ-3 & 0% DSPC & 2.5% PEG | 45：0：52.5：2.5 | 8 | 87.1 | 0.163 | 97.5 |
| 4 | 45% LQ-3 & 0% DSPC & 3% PEG | 45：0：52：3 | 8 | 86.8 | 0.190 | 97.5 |
| 5 | 45% LQ-3 & 0% DSPC & 3.5% PEG | 45：0：51.5：3.5 | 8 | 76.3 | 0.210 | 97.8 |
| 6 | 45% LQ-3 & 0% DSPC & 4% PEG | 45：0：51：4 | 8 | 87.0 | 0.234 | 98.6 |
| 7 | 45% LQ-3 & 0% DSPC & 5% PEG | 45：0：50：5 | 8 | 110.3 | 0.235 | 98.2 |
| 8 | 45% LQ-3 & 2% DSPC & 1.6% PEG | 45：2：51.4：1.6 | 8 | 95.9 | 0.149 | 98.6 |
| 9 | 45% LQ-3 & 2% DSPC & 2% PEG | 45：2：51：2 | 8 | 83.0 | 0.175 | 97.6 |
| 10 | 45% LQ-3 & 2% DSPC & 2.5% PEG | 45：2：50.5：2.5 | 8 | 92.2 | 0.355 | 97.5 |
| 11 | 45% LQ-3 & 2% DSPC & 3% PEG | 45：2：50：3 | 8 | 72.3 | 0.193 | 97.3 |
| 12 | 45% LQ-3 & 2% DSPC & 3.5% PEG | 45：2：49.5：3.5 | 8 | 68.0 | 0.231 | 97.4 |
| 13 | 45% LQ-3 & 2% DSPC & 4% PEG | 45：2：49：4 | 8 | 69.0 | 0.235 | 97.9 |
| 14 | 45% LQ-3 & 2% DSPC & 5% PEG | 45：2：48：5 | 8 | 75.7 | 0.292 | 97.6 |

**Table S4.** Formulations and physicochemical characteristics of LQ-3 iPLX and SM-102 LNP.

| LNP | Group | % mole ratios (ionizable lipids/DSPC/Cholesterol/DMG-PEG2k) | N/P | Isoelectric point | Zeta potential (mV) |
|---|---|---|---|---|---|
| 1 | LQ-3 iPX | 45：0：53：2 | 8 | 6.9 | -2.6 |
| 2 | SM-102 LNP | 50：10：38.5：1.5 | 6 | 6.8 | -4.0 |

**Table S5.** Mean interaction energy between mRNA and either SM-102 or LQ-3.

| Interaction group | Coulomb (kJ/mol) | van der Waals (kJ/mol) | Total (kJ/mol) |
|---|---|---|---|
| mRNA-SM-102 | -135.5 | -457.9 | -593.4 |
| mRNA-LQ-3 | -307.3 | -929.6 | -1236.9 |
